# Task-induced topological and geometrical changes in whole-brain dynamics predict cognitive individual differences

**DOI:** 10.64898/2026.04.19.719533

**Authors:** Ruiqi Chen, Hayoung Song, ShiNung Ching, Todd Braver

## Abstract

Across the last three decades, functional magnetic resonance imaging (fMRI) research – through both resting-state (rsfMRI) and task-based (tfMRI) studies – has greatly advanced our understanding regarding the neural basis of cognition. Yet the mechanistic relationship between rsfMRI and tfMRI is still poorly understood. In particular, it remains unclear how and why the brain activation patterns observed during the resting state are linked to cognitive functioning and individual differences present during task performance. Here, we test a unifying computational account which postulates that task contexts modulate the nonlinear attractor landscape and associated dynamical properties of the brain present under resting conditions, and further that the nature of this modulation is impacted by meaningful cognitive individual differences. To test this account, we develop a joint rsfMRI-tfMRI modeling and analysis framework called Mesoscale Individualized NeuroDynamics with eXogenous inputs (MINDy-X) and apply it to resting and N-back working memory task data from the Human Connectome Project. We first validated that the joint model can simulate and predict both rsfMRI and tfMRI data accurately, consistent with a common underlying dynamical system. Analyses of this joint model revealed that task-related modulation bifurcated the predominantly multistable attractor dynamics present during the resting state towards a predominantly monostable dynamics observed during N-back task states. This topological shift was also accompanied by a geometric reconfiguration, with the task state characterized by an enrichment of dynamical attractor “motifs” clustered around the frontoparietal (FPN) and default mode (DMN) networks. Task-related modulations of this attractor landscape were further subject to clear individual differences, such that individuals who did not exhibit a shift in attractor topology were more error-prone and less cautious in responding, while closer geometric proximity to the FPN and DMN motifs explained additional aspects of task performance. N-back behavior was best characterized by the combination of topological and geometric properties present in both task and rest states, suggesting that they each account for unique aspects of individual variability. The current work supports a novel computational framework for understanding the whole-brain neural activity patterns observed during rsfMRI and tfMRI as reflecting different states within a common non-linear dynamical system. This framework provides a new vocabulary for characterizing cognitive functioning in terms of the unique geometric and topological configuration of the associated attractor landscapes, with the potential for wide application in many domains of basic and clinical neuroscience research.

## 1 Introduction

The last three decades have witnessed amazing growth in cognitive neuroscience research, due in large part to advances in the use of functional magnetic resonance imaging (fMRI) neuroimaging techniques. Historically, task-based fMRI (tfMRI) was the primary approach used to identify neural correlates of cognition, by contrasting activation patterns observed under different task conditions. Later, resting state fMRI (rsfMRI) established a new paradigm from which to identify group-related and individual differences in cognitive functioning in terms of brain organization and dynamics. The rsfMRI paradigm has been enormously successful in advancing understanding within many cognitive domains, and across both healthy and patient populations (Biswal & Uddin, 2025). However, a fundamental question remains unsolved: how and why does the neural dynamics present during the resting state relate to cognitive functioning assessed during task contexts?

Here, we put forward a unifying computational account which postulates the presence of a common mechanism present during both spontaneous resting states and controlled cognitive task states, in which state changes reflect modulations occurring within a single underlying dynamical system. This hypothesis itself is not novel. Indeed, in terms of both mental and neural dynamics, resting and controlled cognitive task states have been frequently suggested to reflect different facets of a continuum, rather than being fundamentally distinct (Christoff et al., 2016). The resting state undoubtedly involves a rich stream of mental activities (Gonzalez-Castillo et al., 2021; Ke et al., 2025), and, correspondingly, the resting brain is well-understood to never truly be at rest (Deco, Jirsa, & McIntosh, 2013). Conversely, off-task mental activities (such as mind wandering) necessarily occur within the context of cognitive task states (Seli et al., 2018). Spontaneous neural fluctuations, while often treated as noise, have also been shown to bias single-trial neural and behavioral responses to task stimuli (Baria et al., 2017; Wu et al., 2024). Therefore, it stands to reason that spontaneous resting and controlled task states might both be supported by the same brain mechanism, which further suggests the predictive utility of resting state dynamics for understanding cognitive functioning, including both group-based and individual differences.

Nevertheless, it has been challenging to even formulate testable predictions regarding this shared mechanism underlying resting state and task-related brain dynamics, let alone to successfully capture it computationally and analytically. This problem occurs because of the lack of a common vocabulary for characterizing resting and task-related activation patterns. Traditionally, rsfMRI and tfMRI data have been analyzed independently, and with distinct methodological approaches. In particular, tfMRI studies are typically oriented towards identifying differences in neural activation patterns associated with experimental contrasts (Friston et al., 1994). In the absence of experimental manipulations, such neural patterns are ill-defined within rsfMRI. Conversely, analysis techniques, such as functional connectivity (FC) and related approaches, are most commonly used in rsfMRI studies, but have also been found to be informative for tfMRI as well (Friston et al., 1997; Gonzalez-Castillo & Bandettini, 2018; Huang et al., 2024). In fact, influential studies have shown that whole-brain FC organization is quite consistent across both resting and task states (Cole et al., 2014; Krienen et al., 2014), which might provide a backbone for the transformation of cognitive representations across brain regions (Cole et al., 2013; Ito et al., 2022), leading to the predictive power of resting state FC for task performance (Tavor et al., 2016). Yet, because the FC measure reflects an emergent statistical summary of co-activation patterns, FC-based models by themselves cannot explain how FC emerges mechanistically, and moreover, why FC might be similar across resting and task states (Friston, 2011). Further, similar to other common fMRI features, FC metrics are not intrinsically sensitive to the temporal dynamics of the data (even though they are commonly treated as if they are), because they are unaffected by shuffling of fMRI volumes in time. Therefore, a mechanistic account for rsfMRI and tfMRI dynamics remains missing.

As an alternative approach, the modeling of fMRI data within the lens of dynamical systems theory has the potential to provide a new vocabulary and set of analytic tools from which to understand resting-state and task-based brain activity patterns in terms of common and unifying mechanisms. Within the dynamical system framework, fMRI timeseries are often assumed to be generated according to dynamical principles that are well-captured by the ordinary differential equation *ẋ* = *f* (*x*) or the discrete-time difference equation *x_n_*_+1_ = *f* (*x_n_*), in which the variable *x* represents the neural state at a specific time. The system evolves according to the direction defined by the vector field *f* (*x*), forming a trajectory within state space. The vector field *f* (*x*) thus implicitly captures the temporal structure of the fMRI timeseries by describing the attractor landscape of the dynamics, where an “attractor” is a state (or a continuum of states) that activation trajectories will tend towards. By integrating these trajectories according to dynamical principles (potentially under noise perturbation), dynamical systems-based computational models can simulate realistic fMRI timeseries that recreate key statistics of the data such as FC (Deco, Ponce-Alvarez, et al., 2013). As such, these computational models reveal the generative mechanisms that give rise to fMRI-measured whole-brain dynamics.

Dynamical systems-based modeling approaches have been successfully applied to both rfMRI and tfMRI in previous work (considered further in the Discussion Section), but to our knowledge, only a few studies have directly investigated and jointly compared both resting and task-based states. Kashyap et al. (2025) fit different models for rsfMRI and each tfMRI dataset within the Human Connectome Project (HCP; Van Essen et al., 2013) separately. They found that single trial reaction time (RT) could be better predicted by a linear combination of both the rsfMRI and tfMRI models rather than each model alone. In contrast, (Englert et al., 2025) constructed a recurrent neural network (RNN) with weights given by resting state FC and inputs being task activation statistical maps. They showed that task-state fMRI evolution aligned with the prediction of the input-driven model, supporting the existence of a common underlying mechanism modulated by tasks. We suggest that neither of these frameworks is optimal. Instead, we argue that, resting and task states should be treated as a continuum, and that the most appropriate analytic framework for joint rsfMRI-tfMRI dynamics is a unified dynamical systems model, in which task effects are implemented as modulatory factors that intrinsically impact the trajectory of activation state dynamics. Moreover, because such modulatory effects are likely to be influenced by relevant cognitive individual differences (Schultz & Cole, 2016), the optimal framework is one that is constructed from individualized models of brain network organization – a feature that has been lacking from all previous studies.

To rigorously test this theoretical account, we developed a joint rsfMRI-tfMRI computational modeling and analysis framework called Mesoscale Individualized NeuroDynamics with eXogenous inputs (MINDy-X). MINDy-X is an adaptation and significant extension of MINDy, a generative framework developed for modeling and analysis of rsfMRI data (Singh, Braver, et al., 2020). Similarly to MINDy, MINDy-X contains hundreds of interconnected neural masses representing brain areas. Previously, the MINDy framework has been shown to fit rsfMRI data more reliably and accurately than alternative linear models, supporting the characterization of rsfMRI data as reflecting a nonlinear dynamical system. Further, we have utilized MINDy to identify and define the attractor landscape present within brain networks during the resting state (Chen et al., 2025), and have also extended the framework to capture attractor dynamics in M/EEG data (Singh et al., 2025). Here, we utilize MINDy-X to leverage its key advance, which is that it enables modeling the effects of task-related variables and how they modulate spontaneous dynamics. This makes it possible to investigate and compare the brain’s dynamical landscape across different cognitive states, as well as across individuals.

We applied MINDy-X to study the geometry and topology of whole-brain dynamical landscapes present across resting states and working memory task contexts, by using fMRI data from the Human Connectome Project (Van Essen et al., 2013). We found that the brain dynamics during both resting state and task state can be well-approximated by the MINDy-X model by including task modulatory factors, supporting the hypothesis of a unifying mechanism across resting and task states. Critically, however, task-related cognitive states were found to be characterized by significant topological and geometrical changes within the whole-brain dynamics. Topologically, these control-demanding cognitive task states reshaped the landscape from that of multistability and/or oscillatory patterns towards monostability. Geometrically, task-state dynamical attractors were characterized by strong frontoparietal network (FPN) activation patterns that were absent in resting state. Further, model-based characterizations of resting state and task state attractor landscapes, involving both the FPN and default mode network (DMN), could predict individual differences in task performance, while the resting and task-based characterizations each explained unique components of individual difference variation. Together, these findings suggest that fMRI-measured resting-state and task-state brain dynamics jointly rely on a unifying mechanism that reflects cognitive individual differences. The MINDy-X framework provides a sufficiently general approach that can enable joint rsfMRI-tfMRI analyses across a wide range of cognitive domains, and a new vocabulary for characterizing the temporal structure of whole-brain activation patterns within a variety of population groups.

## 2 Methods

### 2.1 Data acquisition and preprocessing

All fMRI and behavioral data in this study were originally collected as part of the Human Connectome Project (HCP; Van Essen et al., 2013). The data were publicly available at the HCP project website, after signing the data use terms. For this study, we focused on the fMRI data acquired during resting state and N-back working memory task scans. In the resting state, participants fixated on a bright cross hair overlaid on a dark background, presented in a darkened room. The N-back task contains 0-back and 2-back blocks, each with ten trials of 2.5 seconds. In 0-back blocks, a visual target was presented at the beginning of the block, followed by ten stimulus each presented for two seconds with a 500ms interval. Participants were required to respond when the target is presented. In 2-back blocks, participants were required to respond when the current stimulus is the same as the second last one. There were a total of eight blocks (80 trials) for each condition. For more details, please refer to (Barch et al., 2013).

All fMRI data were collected on a single 3T scanner with a TR of 720ms and 2mm isotropic voxels. Resting state scans contained four runs of 1200 TRs (around 15 minutes), collected during two separate days. On each day, one run had right-to-left (RL) phase encoding and the other had left-to-right (LR) encoding. N-back task scans contained one RL run and one LR run of 405 TRs each (around 5 minutes). We preprocessed rsfMRI and tfMRI data analogously following (Siegel et al., 2017), which were shown to suppress spurious brain-behavior associations due to head motion. We obtained the “minimally preprocessed” fMRI data from HCP (without ICA-FIX correction, as it is less conventional for tfMRI). Then, we de-meaned, standardized, and detrended the fMRI timeseries for each vertex in each run. When de-meaning and standardizing tfMRI data, we used the mean and standard deviation of the corresponding rsfMRI timeseries (with the same phase encoding) instead of its own mean and standard deviation, so that the two types of scans were normalized in a common manner. We then performed a linear regression per vertex to remove the effect of the following confounders (Siegel et al., 2017): (1) 12 rigid-body movement regressors defined by HCP, and their quadratics; (2) the top five principal components of the white matter signal and those of the cerebrospinal fluid signal; and (3) the mean signal from the gray matter. Further, we applied “motion scrubbing” (Power et al., 2012) to linearly interpolate volumes with high framewise displacement (FD), defined as FD above 0.2mm for rsfMRI and 0.9mm for tfMRI (Etzel, 2023). We excluded all participants with missing runs or runs with more than 1/3 high-FD volumes, which resulted in a subset of 537 participants remaining for the main analysis. After motion scrubbing, the vertex timeseries were averaged according to the Schaefer atlas with 200 parcels (Schaefer et al., 2018). Then, we de-meaned and standardized the parcel timeseries. For tfMRI, we used the mean and standard deviation during the block intervals to normalize the data, such that the data would be zero-meaned with unit variance during block intervals, similar to that in resting state. As the final step of preprocessing, we deconvolved the BOLD data to obtain the underlying neural activities. This step will be further described and explained in Section 2.3.

### 2.2 Unifying model for resting and task states

In this study, we assumed that the fMRI-measured ultra-slow whole-brain dynamics during resting state and cognitive task states are generated by the same underlying mechanism. We aimed to show that this mechanism can be modeled as a dynamical system driven by effective connections between brain regions, with the activation levels of these regions modulated by cognitive processing.

Mathematically, we assumed the fMRI signal was generated by a discrete-time dynamical system consisting of *n* interconnected brain regions, each modeled by a nonlinear neural mass. The dynamics of the system are governed by the following difference equation:

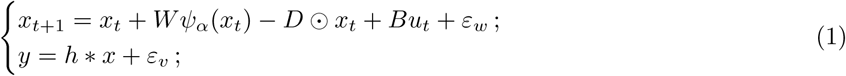

where *x_t_* ∈ R*^n^* represents the overall activation level (hidden state) of each brain region at volume *t* and *y_t_* ∈ R*^n^* represents the observed fMRI BOLD signal. We assumed that the neurovascular coupling can be modeled as a linear system with impulse response *h* (the haemodynamic response function, HRF), as in traditional GLM-based fMRI analysis (Friston et al., 1998). Under this assumption, the BOLD signal is generated by convolving (indicated by ∗) the neural activity *x* with the impulse response *h*, under the influence of the *observation noise ε_v_*.

The dynamics of the neural hidden state *x* are driven by four different sources: (1) the recurrent input from other regions *Wψ_α_*(*x_t_*), where *W* ∈ R*^n^*^×^*^n^* represents the *effective connectivity* between regions and *ψ_α_*(*x_t_*) ∈ R*^n^* represents the *output* of each region (detailed below); (2) the decay −*D* ⊙ *x_t_*that reflects local inhibition within each region, with *D* ∈ (R^+^)*^n^* referred to as the *decay rate* and ⊙ indicating the element-wise product; (3) the task-related modulations *Bu_t_*, where *u_t_* ∈ R*^n^* represents any task-related variables and *B* ∈ R*^n^*^×^*^k^* models the coupling between task variables and brain activation level; and (4) the *process noise ε_w_* intrinsic to the system. Both the process noise *ε_w_* and the observation noise *ε_v_* are assumed to be zero-meaned, Gaussian, and uncorrelated across regions. The architecture can be considered as a “vanilla” or Amari-type recurrent neural network (RNN; Amari, 1977). However, instead of a fixed-width tanh activation function, MINDy-X uses a parameterized sigmoidal function that allows each region to respond in a more or less step-like fashion:

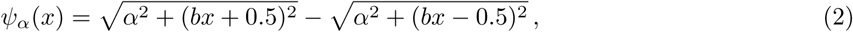

where *b* = 20*/*3 is a fixed scaling factor and *α* ∈ R*^n^* controls the slope/curvature of the response function for each region (see Figure 1C). All operations in (2) are element-wise. *α* can be considered as reflecting the variability of neuronal activities within each region (Marreiros et al., 2008). A small *α* corresponds to low neuronal variability and a more step-like regional response, while a large *α* corresponds to high variability and a flatter response profile.

**Figure 1:**
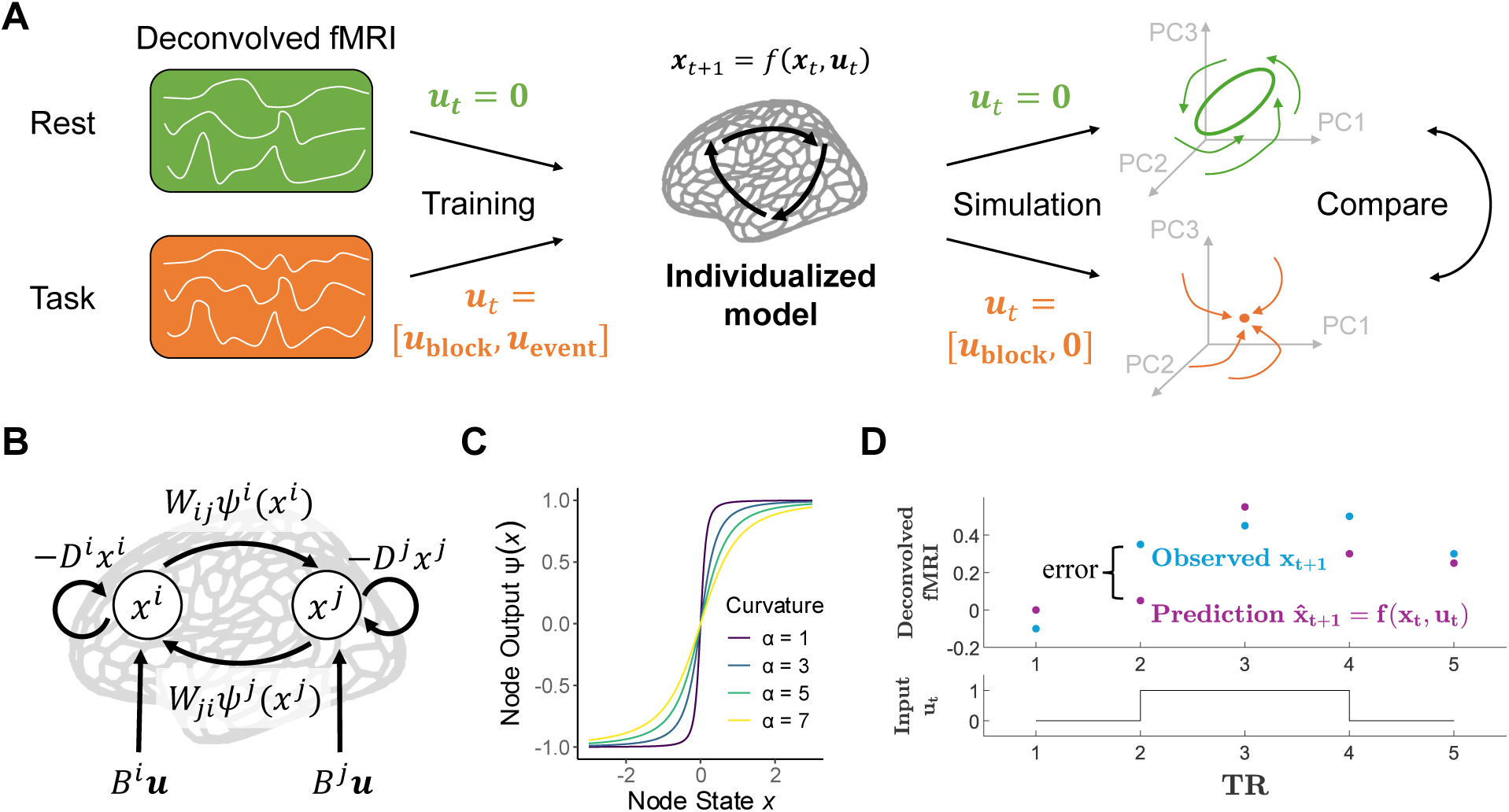
MINDy-X schematics. (A) Joint rsfMRI-tfMRI analysis with MINDy-X. A MINDy-X model is fit on one participant’s deconvolved rsfMRI and tfMRI data to predict the next volume *x_t_*_+1_ based on the current volume *x_t_* and task-relevant inputs *u_t_*. After fitting, the model is simulated with and without inputs to reveal the attractor landscape of brain dynamics in each task and resting conditions. (B) MINDy-X architecture. In this study, each MINDy-X model contains 200 neural masses representing 200 cortical regions in the Schaefer atlas (Schaefer et al., 2018). Activation of each region is driven by recurrent inputs *Wψ_α_*(*x*) from other regions, task-related modulations *Bu*, and local inhibition (decay) −*Dx*. (C) Activation function *ψ_α_*(*x*) as the reciprocal of slope *α* changes. Typical fit of *α* was around 5. (D) Illustration of MINDy-X fitting process. The model predicts the next deconvolved neural state based on the current state *x_t_* and task variables *u_t_*. The parameters were optimized to minimize the error between the prediction *f* (*x_t_, u_t_*) and actual state *x_t_*_+1_.

To establish the relationship between cognition and neural dynamics, the key component of the MINDy-X model is the task-related term *Bu_t_*, interpreted as a modulation to the neural activation level. In this paper, we simply designed *u_t_* ∈ R^2^ to be two box-car regressors indicating task conditions:

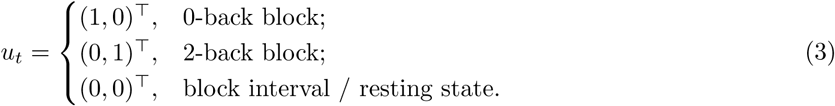

Accordingly, *B* ∈ R*^n^*^×2^ contained two columns representing how the excitability of brain regions deviated from resting state during 0-back and 2-back blocks, respectively. Importantly, such deviation is expected to induce a change in the geometry and possibly topology of the dynamical landscape, revealing how cognitive processes influence the spatiotemporal structure of brain dynamics. The task state block intervals were treated analogously to the resting state in the current paper, but in principle, it could also be modeled as a separate condition by including a third box-car regressor.

Moreover, while we adopted a relatively simple task effect model here, MINDy-X can incorporate any effects representable by a GLM. For example, to model the lasting effects of an event-related variable on the dynamics, one can include time-lagged copies of the variable in *u_t_* and treat *B* as a linear convolution kernel. While block-type variables can be interpreted as a persistent *modulation* of the dynamical landscape to suit task demand, event-related variables can naturally be interpreted as a transient *drive* that pushes the system to computationally relevant states (Song et al., 2025). In other prior work, we pursued a related, but distinct approach, in using MINDy estimates of the resting-state dynamics as a pre-processing step that could then be filtered out of task contexts, to better estimate task-related modulations on the system without the contribution of intrinsic dynamics overlaid (Singh, Wang, et al., 2022).

### 2.3 Dual estimation of states and parameters

Because only the output *y_t_* of the model was observed, both the parameters (*W, α, D, B*) and the hidden states (*x_t_*) of the model need to be estimated in an interdependent manner, leading to a challenging dual estimation problem (Singh, Wang, et al., 2020). To simplify this problem, we made two assumptions. First, we assume that the HRF *h* follows a canonical form defined in (Friston et al., 1998). While in reality, the HRF might vary across regions and individuals (Aguirre et al., 1998), this assumption is nonetheless conventional in GLM-based tfMRI analysis. Second, according to common estimations (Buxton, 2013), we assume that the ratio between the variance of the signal *x_t_* and that of the observation noise *ε_v_* is around 50:1 across all frequency bands. Based on these two assumptions, the maximum likelihood (minimum squared error) estimate of the signal *x_t_* can be obtained through the following Wiener deconvolution, represented in frequency domain:

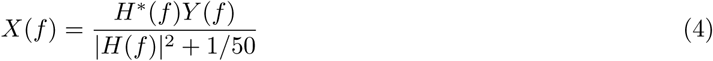

where *X*(*f* ), *Y* (*f* ) and *H*(*f* ) are the Fourier transform of *x_t_*, *y_t_* and *h_t_* respectively. As a last step of our preprocessing pipeline, we performed this deconvolution to obtain the “neural” timeseries *x_t_*, which was used for modeling and all subsequent analysis.

Given the neural timeseries *x_t_*, we tried to optimize the model parameters to minimize the error between the predicted and observed value for the next state *x_t_*_+1_. To prevent overfitting and improve interpretability, we regularized the connectivity matrix *W* to enforce a “low rank plus sparse” structure. Specifically, *W* is parameterized as the sum *W* = *W_s_* + *W*_1_*W*_2_^⊤^, where *W*_1_*, W*_2_ ∈ R*^n^*^×^*^k^* with *k* much smaller than the full dimension *n* (in this study, *n* = 200 and *k* = 72). We also regularize via the *ℓ*_1_ norm (absolute sum of entries) of the matrices *W_s_*, *W*_1_ and *W*_2_ to promote a sparse solution (Donoho, 2006). Moreover, we place an extra *ℓ*_1_ regularization over the diagonal elements of *W_s_* to suppress its redundancy with the decay term. The loss function is thus

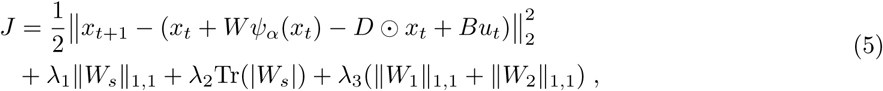

where ∥*A*∥_1,1_ := Σ*_i,j_* |*A_ij_*| denotes the *ℓ*_1_ norm and Tr(|*A*|) := Σ*_i_* |*A_ii_*| denotes the absolute sum of diagonal elements. We set the regularization weights as *λ*_1_ = 0.075, *λ*_2_ = 0.2 and *λ*_3_ = 0.05 following (Chen et al., 2025).

Note that the MINDy-X loss function is nonlinear with respect to not only the states *x_t_*but also the parameter *α*, so it cannot be transformed into a linear regression problem as in NARMAX models (Billings, 2013), such as Kashyap et al. (2025). To minimize this loss, we used a gradient descent algorithm with Nesterov Accelerated Adaptive Moment Estimation (NADAM; Dozat, 2016). During each minibatch, we randomly sampled 300 triplets of (*x_t_, u_t_, x_t_*_+1_) and updated the parameters according to the average loss. The samples were drawn uniformly across all scanning runs (both resting state and task scans), with *x_t_* and *x_t_*_+1_ being consecutive volumes from the same run. We trained the models for 5000 minibatches as the test-retest reliability of parameter started to drop after that. Following the gradient descent, we performed a global rescaling of the parameters to compensate for regularization bias. We fit three scalar parameters *p_W_ , p_D_ , p_B_* ∈ R to minimize the prediction error 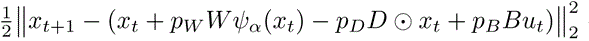 over all training data using linear regression, and factored them into *W* , *D* and *B*. For validation purposes (Section 3.1), we trained the models using each participant’s data during one visit and tested on the other visit, using the HCP test-retest dataset. For general analyses, we trained one model for each participant using all preprocessed resting state and N-back task fMRI data in the HCP young adult dataset. It is worth noting that each MINDy-X model can be trained within 15 seconds on a standard computer, suggesting that the approach is highly scalable, even for very large datasets such as ABCD (Casey et al., 2018), Cam-CAN (Taylor et al., 2017), and UK Biobank (Miller et al., 2016).

### 2.4 Alternative models

We compared the goodness-of-fit and parameter test-retest reliability of MINDy-X model against several linear models, with decreasing complexity: (1) “Brainwise linear”: a linear dynamical model with effective connectivity between regions. This model can be considered as replacing the nonlinear activation function *ψ_α_*(*x*) in MINDy-X with the identity function. It can also be viewed as the vector autoregression model VARX(1, 1): *x_t_*_+1_ = *Ax_t_* + *Bu_t_* + *ε_t_*_+1_. (2) “Decay only”: a linear dynamical model with parcelwise decay but no global connectivity, namely the scalar autoregression model ARX(1, 1): *x_t_*_+1_ = *D* ⊙ *x_t_* + *Bu_t_* + *ε_t_*_+1_; (3) “GLM”: a pure noise process model, namely the general linear model GLM: *x_t_* = *µ* + *βu_t_* + *ε_t_*. To prevent overfitting, we applied an *ℓ*_1_ regularization on the *A* matrix for the brainwise linear model, with regularization weight *λ* ranging from 0, 0.001, 0.005, 0.01 to 0.05. We selected the solutions with *λ* = 0.01 based on cross-validated fit and parameter reliability. All models were fit on the same deconvolved fMRI data and cross-validated between the two visits (waves) of each participant, as MINDy-X did. The brainwise linear models were fit using glmnet::glmnet in R. The decay only and GLM models were fit using simple linear regression in MATLAB.

### 2.5 Model simulation and attractor identification

In Section 3.1.2, we aimed to verify MINDy-X’s ability to simulate neural time series that recapitulate key statistics of observed fMRI data, given appropriate input and noise structure. Starting from a random initial condition *x*_0_ ∈ R*^n^* (with entries drawn independently from the standard normal distribution), we first let the models run freely for 1000 time steps to converge to its steady-state before introducing task inputs. In each step, Gaussian white noise was injected independently to each parcel, with zero mean and variance equal to the mean squared prediction error for that parcel on the validation set. After the 1000-step warm-up, we provided the model with the same input sequence *u_t_* as in the training data, in addition to the noise. We thus obtained a sequence of the hidden state data *x_t_* of the same length as *u_t_*, which was then compared with the observed data in terms of functional connectivity (FC) and GLM-estimated effects of task variables.

While the noise-driven MINDy-X simulation generates realistic neural traces, a noise-free simulation provides a clearer picture of the mechanism underlying observed dynamics. Most importantly, noise-free simulations can reveal the attractors of the system, i.e., states (or continua of states) that the system naturally evolves towards. We adopted the same method as in Chen et al. (2025) to identify the attractors. To simulate the dynamics during resting-state/0-back/2-back condition, we fixed the input *u_t_* ≡ *u* to the value corresponds to that condition defined in Section 2.2. Starting from 120 random initial conditions, we forward iterated the model for 1600 steps under the input without adding noise. For some models containing extremely slow regions, we extended the simulation until trajectories converged. We treated a simulated trajectory as already converged to a stable equilibrium (fixed point) if 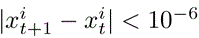 for every parcel *i* and every time point *T* − 10 ≤ *t* ≤ *T* − 1, where *T* is the number of simulated time steps. We clustered the final states *x_T_* of all trajectories that converged to equilibria based on a Euclidean distance threshold of 0.1, and selected the cluster centroids as candidates of stable equilibria. Other than equilibria, many simulated trajectories converged to a stable limit cycle (a sustained oscillation). We identified such trajectories by the criterion that it had entered and then left the neighborhood {*x* ∈ R*^n^* | ∥*x* − *x_T_* ∥_2_ *<* 0.5} at least once. The interval during the last entry to this neighborhood and the end of simulation was chosen as the period of the limit cycle, and the trajectory during this period was extracted to represent the limit cycle. We also combined duplicated limit cycles based on a distance threshold. We verified the identified attractors by visually inspecting the trajectories after dimensionality reduction (e.g., Figure 5A), and removed 14 out of 537 models where the numerical procedure might have failed.

Notably, we found at least one attractor in all models across all conditions, and we did not observe metastable or chaotic behavior. We thus categorized all models into three types within each condition: (1) monostable type, with only one globally stable equilibrium, and no stable limit cycles; (2) multistable type, with more than one stable equilibria and no stable limit cycles; and (3) oscillatory type, with at least one stable limit cycle. Such categorization was based on the topology of the vector field governing the model dynamics, regardless of geometrical variations (e.g., the exact location of the equilibria, or the period of the limit cycles). Importantly, when driven by mild noise, these three types of systems showed qualitatively different long-term behavior: the monostable system fluctuated around a single stable state, while the multistable system could fluctuate between several stable states intermittently. The oscillatory system, on the contrary, showed sustained noisy oscillations. Therefore, these three types of dynamics can be treated as three distinct generative mechanisms, which could then be potentially linked to cognitive individual differences, as part of our following analyses.

### 2.6 Geometrical characterization of whole-brain dynamics

The whole-brain dynamics captured by MINDy-X models differed across individuals and cognitive states not only in topology but also in geometry. We were primarily interested in the distribution of stable equilibria over the state space, as well as in the relationship between the empirical neural states and the theoretical (model-based) equilibria. In Chen et al. (2025), we showed that during resting states, these equilibria formed several clusters that could be associated with different functional brain network activation patterns. We performed a similar clustering analysis in the current paper for each cognitive condition (rest/0-back/2-back) separately. The stable equilibria (represented by *n*-dimensional vectors) from all models under the corresponding input were clustered using K-means algorithm, with the number of clusters *K* ranging from two to ten. We selected the optimal *K* as the local minimum of clustering instability index (Lange et al., 2004), which turned out to be four for resting state and 0-back conditions, and three for the 2-back condition. To assess the quality of the clustering solution, in Figure 6, we projected all stable equilibria to the first two “principal gradients” of FC defined in Margulies et al. (2016). We also visualized the cortical activation patterns associated with the cluster centroids, and named the clusters accordingly. In the resting state, the four clusters were named “DMN” (default mode network), “VIS” (visual network), “TPN” (task-positive network), and “SAL” (salience network). In the 2-back task state, the three clusters were named DMN, “FPN” (frontoparietal control network) and TPN.

Next, we aimed to summarize individual differences among HCP participants’ whole-brain dynamics, based on the geometry of the identified attractor landscape. We did so by calculating the mean Euclidean distance between the observed neural states (deconvolved fMRI data) *x_t_* ∈ R*^n^* and the model-based attractor centroids *c_k_* ∈ R*^n^* (*k* = 1, 2*, . . . , K*) for each participant during each condition separately. We also performed another analysis akin to that of a Markovian jump process. We assumed that the brain state *x_t_* is switching between *K* meta-states represented by the cluster centroids. We simply categorized each volume *x_t_* into the meta-states with highest cosine similarity to *x_t_*, then calculated the proportion of time each participant spent in each meta-state in each condition. These model-based geometrical features were then used to predict cognitive individual differences in the following analyses.

### 2.7 Predicting task performance with model-based dynamical features

We performed two different types of analyses to establish the connection between model-based features of neural dynamics and individual differences in task performance. The first one was an extreme group analysis. We selected the top 25% participants on 2-back task (in terms of accuracy) into a “high-performing” group and the bottom 25% into a “low-performing” group, after excluding those with an accuracy below 0.6 in any task. We compared the mean distance to centroids and meta-state occupancy between the two groups using MANOVA and post-hoc t-tests.

Beyond the extreme group analysis, we also performed a formal (continuous) analysis using GLM methods. We predicted each participant’s accuracy and log-transformed reaction time on 2-back task using the features introduced before: (1) Bifurc (binary): whether model dynamics bifurcated from rest to task; (2) FPN_time and TPN_time (continuous): mean proportion of time in FPN and TPN meta-states in the 2-back condition (DMN_time was not included as it is collinear with the other two); and (3) dDMN, dFPN, and dTPN (continuous): mean distance between observed neural state and DMN/FPN/TPN cluster centroids in 2-back condition. As an exploratory analysis, we considered several models that were of particular theoretical interest: (1) “Bifurcation”: target ∼ Bifurc; (2) “Occupancy”: target ∼ FPN time + TPN time; (3) “Distance”: target ∼ dDMN + dFPN + dTPN; (4) “Additive”: target ∼ Bifurc + dDMN + dFPN + dTPN; and (5) “Interaction”: target ∼ Bifurc * (dDMN + dFPN + dTPN), which includes the additive model plus the interaction between Bifurc and each distance terms.

Finally, we analyzed whether using the rsfMRI data alone could also predict participant performance during the N-back task; and if so, whether the rsfMRI features explained unique or overlapping variance with the tfMRI features. To eliminate information leakage between sessions, we first re-trained a set of MINDy models using the same parameters but only the rsfMRI data, and repeated all model-based analysis to obtain the resting state attractor centroids (although using the MINDy-X joint models resulted in almost identical results). We selected 508 participants whose (joint and rsfMRI-only) models were free from numerical issues. Then, we calculated the mean distances between rsfMRI neural states and attractor centroids to construct the “Rest-only” model: target ∼ dDMN rest + dVIS rest + dTPN rest + dSAL rest, along with re-estimated “Task-only” model: target ∼ dDMN 2bk + dFPN 2bk + dTPN 2bk. We compared these two models against the joint model that included all seven of these predictors to determine the proportion of unique variance each predicted.

## 3 Results

### 3.1 Unifying dynamics across rest and task states

In this section, we demonstrate that fMRI-measured whole-brain dynamics during both the resting state and N-back task state can be captured by a single unifying nonlinear dynamical model. To this end, we fit and tested MINDy-X models on individual rsfMRI and N-back tfMRI data in the Human Connectome Project (HCP; Van Essen et al., 2013) test-retest dataset with 44 participants (see Section 2.2). Each model is fit to both the rsfMRI and the tfMRI data of one individual in one visit, resulting in 88 models. The models were then tested on the same participant’s rsfMRI and tfMRI data during the other visit. We first verify that the model accurately and reliably predicted the fMRI dynamics across rest and task states, outperforming several linear models. Then, we show that the model parameters are individualized and test-retest reliable. Finally, and most importantly, we demonstrate that the noise-driven simulation of fitted MINDy-X models recreates key statistics of the empirical dynamics, such as functional connectivity (FC) and GLM-estimated neural responses to task. The validated procedure was then applied to the HCP main dataset (*n* = 511 after exclusion, see Section 2.1) to obtain models for further analysis.

#### 3.1.1 Accurate and reliable estimates of dynamics

Here, we aimed to validate the goodness-of-fit and reliability of our model across the HCP resting state and N-back working memory task fMRI data. The task effects were simply modeled by two box-car regressors corresponding to 0-back and 2-back blocks.

We compared MINDy-X against three other kinds of models with decreasing complexity: (1) a linear dynamical model with brainwise connectivity (“Brainwise linear”), namely the vector autoregression model VARX(1, 1): *x_t_*_+1_ = *Ax_t_* + *Bu_t_* + *ε_t_*_+1_; (2) a linear dynamical model with parcelwise decay but no global connectivity (“Decay only”), namely the scalar autoregression model ARX(1, 1): *x_t_*_+1_ = *D* ⊙ *x_t_* + *Bu_t_* + *ε_t_*_+1_; (3) a pure noise process model (“GLM”), namely the general linear model GLM: *x_t_* = *µ* + *βu_t_* + *ε_t_*. All models were fit on the same deconvolved fMRI data and cross-validated between the two visits (waves) of each participant.

We calculated the cross-validated coefficient of determination (R squared) between the actual and predicted next state (*x_t_*_+1_ and *x*^*_t_*_+1_) for all dynamical models (MINDy-X, Brainwise linear, and Decay only), as shown in Figure 2A. All models explained a substantial proportion of variance in *x_t_*_+1_ during both resting state (MINDy-X: *R*^2^ = 0.635 ± 0.004, Brainwise linear: *R*^2^ = 0.632 ± 0.004, Decay only: *R*^2^ = 0.632 ± 0.004) and task scans (MINDy-X: *R*^2^ = 0.576 ± 0.002; Brainwise linear: *R*^2^ = 0.571 ± 0.003; Decay only: *R*^2^ = 0.575 ± 0.002). We performed a repeated-measures ANOVA with model types and testing sessions (rest or task scans) as factors. Both the main effect of model type (*F* (2, 86) = 15.86, *p <* 0.001), main effect of testing session (*F* (1, 43) = 148.4, *p <* 0.001), and the interactions were significant (*F* (2, 86) = 21.34, *p <* 0.001). Dunnett’s test revealed that MINDy-X models significantly outperformed the alternatives on both resting state (MINDy-X vs. Brainwise linear: *t*(111) = 3.418, *p* = 0.002; MINDy-X vs. Decay only: *t*(111) = 3.007, *p* = 0.006) and task scans (MINDy-X vs. Brainwise linear: *t*(111) = 7.090, *p <* 0.001; MINDy-X vs. Decay only: *t*(111) = 2.294, *p* = 0.045). Also, interestingly, the brainwise linear models were less accurate than the decay-only models in some participants, even though in principle they can degenerate into a decay-only model. This suggests that a simple *ℓ*_1_ regularization might be insufficient to prevent overfitting for vectorized linear models (Nozari et al., 2023). On the contrary, the MINDy-X model architecture and regularization techniques provided a good description of individual brain dynamics that were generalizable across different time points.

**Figure 2:**
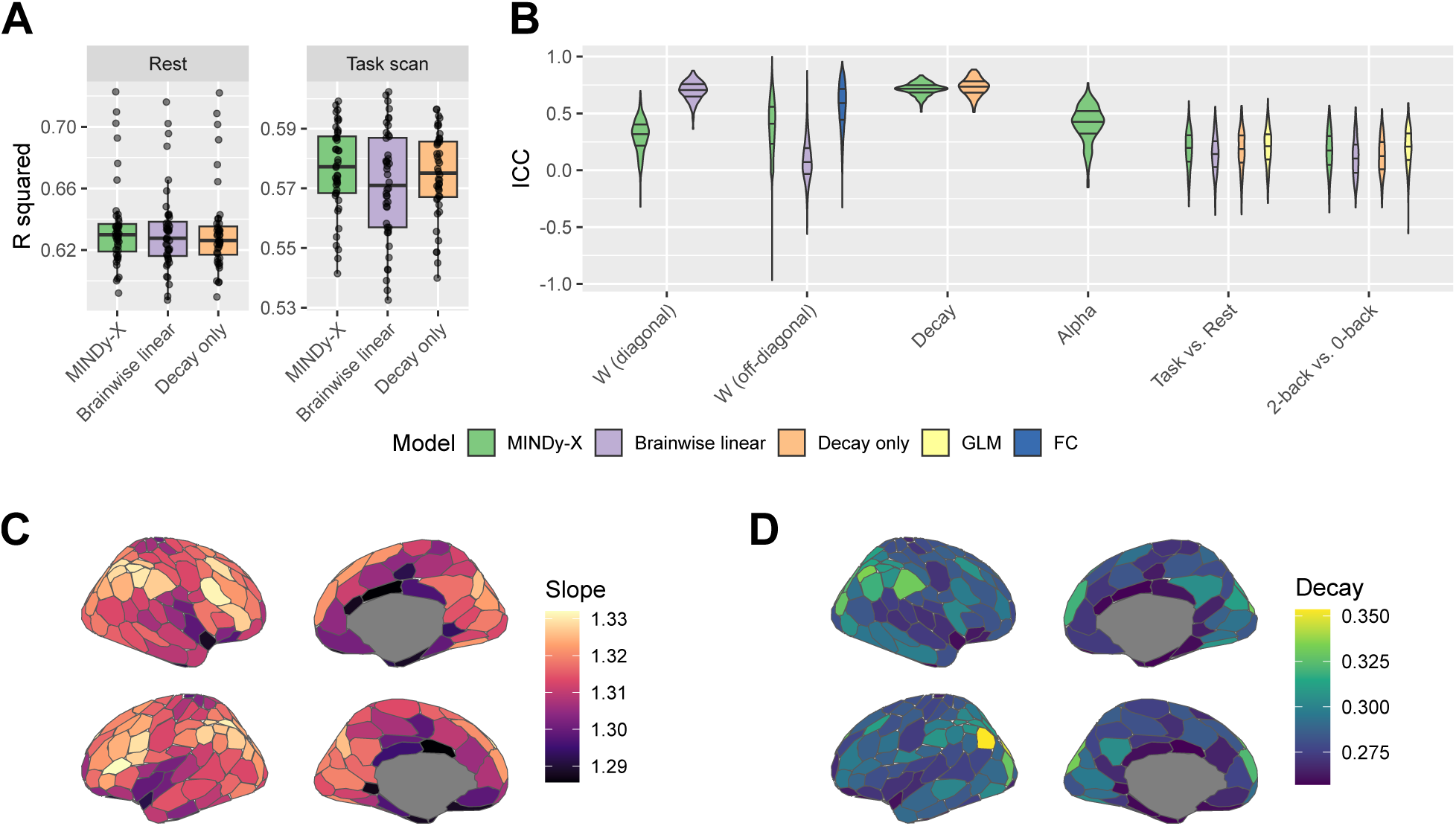
MINDy-X fits were accurate and reliable. (A-B) Goodness-of-fit and parameter reliability on the HCP test-retest dataset. (A) Cross-validated R squared for the prediction of next neural state in resting state and task scans across individualized models. Each dot represents the mean across two models (visits) for on participant. Box plot showed the quartiles of distributions for each model. (B) Intraclass correlation coefficient (ICC) for each model parameter, grouped by parameter types. Models are represented by different colors. ICC for entries of the FC matrix is also shown in blue. Violin plots visualized the distribution and quartiles of ICC over parameters in each group. (C-D) Populational average of model parameters over the HCP main dataset. (C) Derivative of the activation function 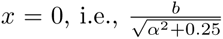 (D) Decay *D* for each region.

**Figure 3:**
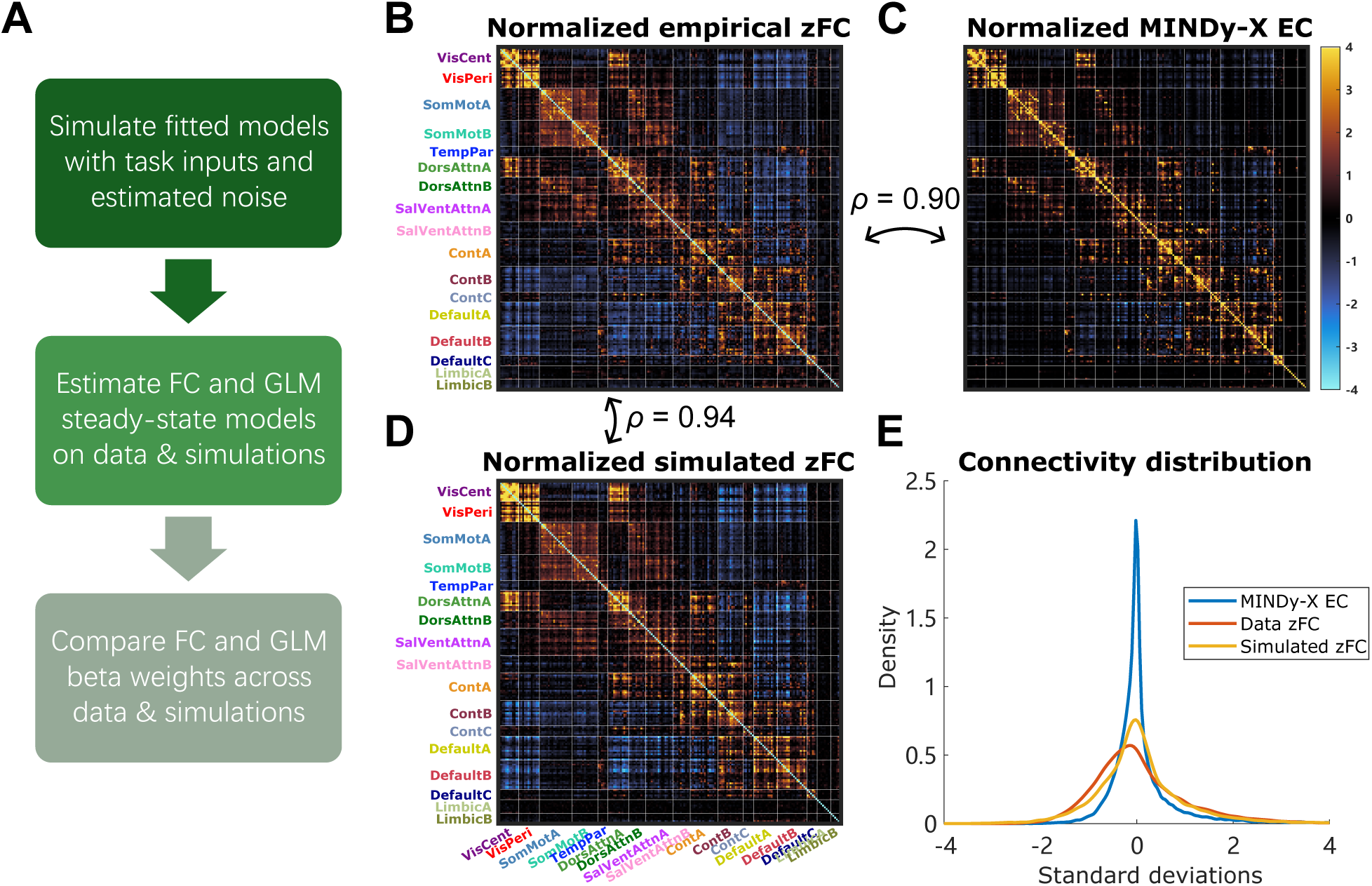
Noise-driven simulation of MINDy-X recreated key statistics of data. (A) Pipeline of the analysis. (B-D) Empirical z-transformed FC, populational averaged MINDy effective connectivity (EC), and z-transformed simulated FC, respectively. Each matrix was normalized to have unit variance (but not de-meaned). Color map was shared across the three panels. Dimensions (parcels) were organized according to the 17 network Schaefer atlas, as indicated by the tick labels, separated by the white lines. (E) Density of the connectivity weights across (B-D).

Next, we analyzed the test-retest reliability of the model parameters to see if the models can capture individual differences in brain dynamics. We computed the intraclass correlation coefficient (ICC) for each parameter in all four types of models, as well as the ICC for the difference between the input weights (for GLM, the regression coefficients) corresponding to 2-back and 0-back blocks. We also calculated the ICC for each entry of the functional connectivity (FC) matrix (Pearson correlation coefficient). The distribution of ICC within each parameter type was visualized in Figure 2B. Consistent with (Shinn et al., 2023), the decay parameters (reflecting the temporal autocorrelation) were extremely reliable in all models. Note that the diagonal elements of connectivity matrix in Brainwise linear models can also be viewed as a decay term, which were also found to be quite reliable. However, the non-diagonal elements of the connectivity matrix were much less reliable in Brainwise linear models, while these elements were still quite reliable in MINDy-X. For task effect estimates, MINDy-X parameters had a similar reliability with traditional GLM regression slopes.

Finally, we moved from the test-retest subset of the data to the full HCP dataset in order to plot the anatomical profile of key MINDy parameters. These anatomical profiles are visualized in Figure 2C-D and Figure 4B-D. Interestingly, the anatomical distribution of the slope parameter aligns with the FPN and DMN, with more step-like slopes present in both lateral and medial frontal and parietal regions. The decay parameter exhibited a different profile with higher decay rates primarily in medial and lateral parietal regions.

**Figure 4:**
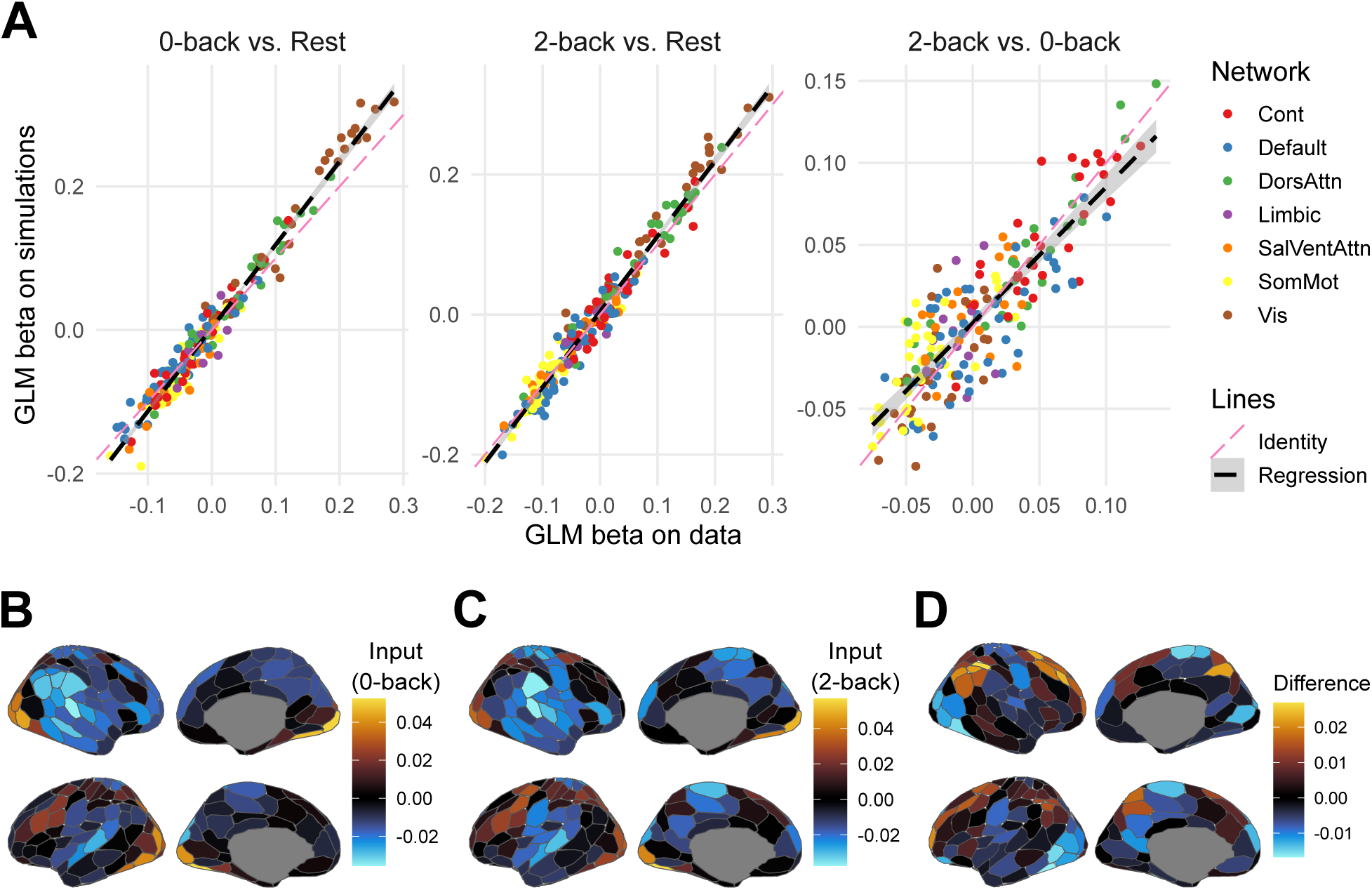
Task effect estimates from MINDy models. (A) Populational averaged beta slopes of GLM models fit on empirical and simulated data for HCP test-retest participants. Each dot represents one parcel and color indicates network assignment in the 7 network Schaefer atlas. Black dashed line represents regression line with shade indicating standard error. Pink dashed line represents the identity line. (B) Input *Bu* for the 0-back condition, i.e., the first column of *B*, averaged across HCP main dataset. (C) Input for the 2-back condition, i.e., the second column of *B*. (D) Difference between (B) and (C).

**Figure 5:**
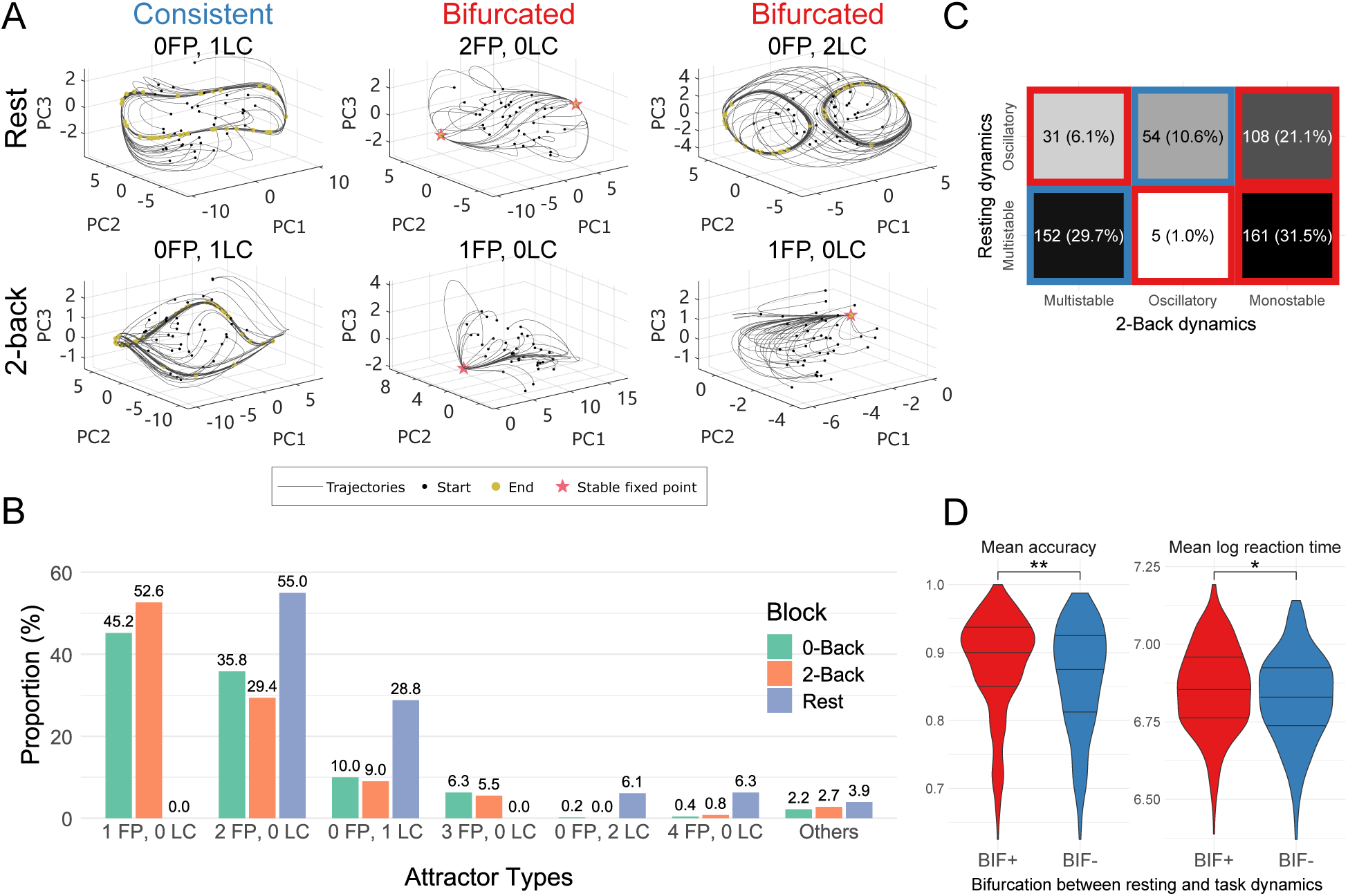
Cautious behavioral strategy was associated with bifurcation between rest and task state neural dynamics. (A) Simulated trajectories and identified attractors of three example models under resting state or 2-back inputs, projected to the first three principal components (PCs) of the trajectories. Black and yellow symbols marked the starting and end points of the trajectories. The end points typically converged to stable equilibria/fixed points (FP) or limit cycles (LC). Number of (stable) FPs and LCs were indicated in the title of each panel. Models showing the same number of FPs and/or LCs in resting versus 2-back conditions were called “consistent” compared to “bifurcated”. (B) Taxonomy of model dynamics across conditions and individuals, categorized according to the number of stable FPs and LCs. Categories with popularity less than five percent in all conditions were grouped into “others”. (C) Contingency table for model types across resting state and 2-back condition. Red and blue blocks correspond to bifurcated and consistent models, respectively. (D) Accuracy and log reaction time in 2-back condition of BIF+ participants, who exhibited a bifurcation in their model from rest-to-task, relative to BIF- who had consistent dynamics, and thus did did not exhibit a bifurcation rest and task states. Statistical significance was indicated by stars (*: *<* 0.05; **: *<* 0.01).

**Figure 6:**
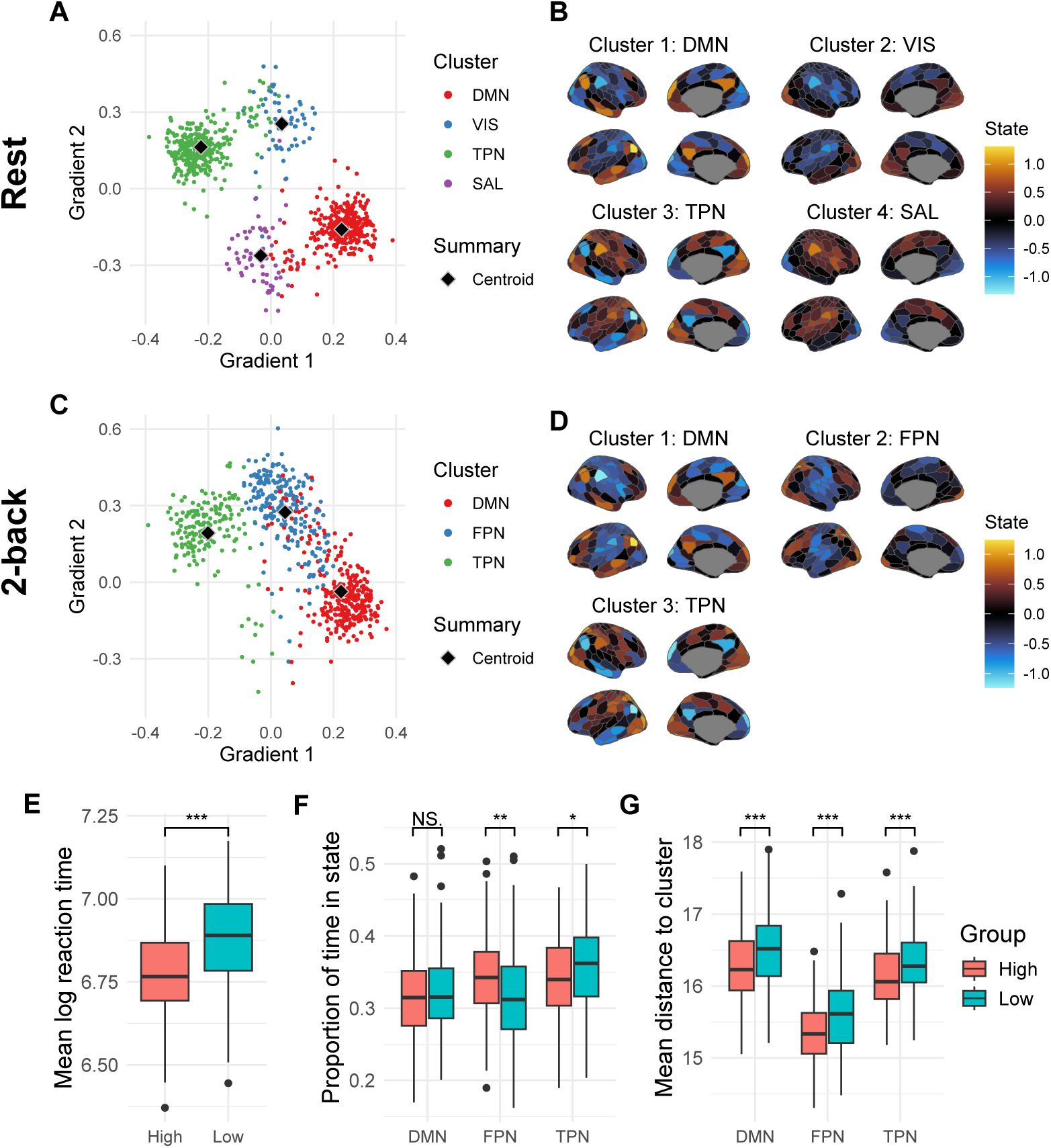
Proximity to attractor motifs predicts working memory performance. (A) Distribution and clustering of stable equilibria in resting state models. The equilibria (200-dimensional vectors) were projected to the first two principal components of FC gradients as defined in Margulies et al. (2016). Cluster membership were indicated by different colors, with centroids marked by black diamond symbols. (B) Cluster centroids (motifs), named according to most prominent network activations. Each centroid was visualized as an activation pattern over the cortex. (C-D) similar to (A-B) but for 2-back condition instead. Axis limits in (C) are the same as in (A). (E-G) Comparison between participants with top and last 25% ranked accuracy. Box plots showed the quartiles, outliers, and non-outlier (within 1.5 IQR from the median) maximum/minimum across individuals. Statistical significance was indicated by stars (NS.: not significant; *: *<* 0.05; **: *<* 0.01, ***: *<* 0.001). (E) Comparison of mean log reaction time between groups. (F) Proportion of time each participant spent in each meta-state in 2-back condition. (G) Mean distance between empirical neural state and attractor cluster motifs within 2-back condition.

Overall, these validation analyses suggest that resting state and N-back task states can be modeled by a single unifying dynamical mechanism, which we found here to be reliable, non-linear and individualized.

#### 3.1.2 MINDy-X recreates connectivity and task effect statistics

We then explored whether MINDy-X can generate neural activation timeseries that recapitulated key statistics of the observed data, including functional connectivity (FC) and N-back task effects estimated by GLM. We forward integrated (1) from random initial conditions to simulate surrogate neural timeseries. To mimic the biological noise inside the brain, we injected Gaussian white noise independently to each parcel during the integration. We assume that the magnitude of prediction error of the model during testing reflects the magnitude of such inherent noise, so the variance of the noise was selected as the mean squared prediction error. We provided the model with the same input sequence *u_t_* as in the training data and obtained the corresponding hidden state trajectory *x_t_*. We then repeated the functional connectivity (FC) and GLM analyses on *x_t_*as in the experimental data (Figure 3A).

First, we examined whether MINDy-X could recreate the observed FC patterns. Figure 3B-D visualize the z-transformed FC (zFC) matrix, the MINDy-X effective connectivity (EC) matrix *W* , and the z-transformed FC of MINDy-X simulations, respectively. The matrices were averaged across participants (and test/retest visits), then normalized to have unit variance (but not de-meaned). While the MINDy-X effective connectivity matrix was already highly correlated with zFC (Pearson’s *ρ* = 0.90), the simulated zFC matched the empirical zFC even better (Pearson’s *ρ* = 0.94). Most remarkably, even though the MINDy-X EC matrix was extremely sparse (excess Kurtosis *κ* = 29.03), the simulated zFC matrix showed a much denser distribution (excess Kurtosis *κ* = 5.98) that matched the empirical data (excess Kurtosis *κ* = 3.14) quite well (Figure 3E), suggesting that the observed FC could be generated by a much sparser scaffold of intrinsic effective connections.

We then analyzed whether MINDy-X simulations resulted in similar GLM-estimated task effects with the actual data. Here the task effects were estimated by the GLM *x_t_* = *µ* + *βu_t_* + *ε_t_* where *u_t_* consisted of 0-back and 2-back block boxcar regressors, as described in previous section. In fact, the anatomical pattern of the MINDy-X input matrix *B* was already quite similar to the canonical N-back activation pattern observed in the GLM *β* coefficients estimated on the data, with prominent activation in lateral frontal and parietal regions of the FPN and dorsal attention networks, and deactivation within DMN regions, albeit with MINDy-X estimated parameters compressed in scale relative to standard *β* coefficients (Figure 4B-D). To examine whether such scale distinction vanished in the GLM analysis, we fit another GLM on the MINDy-X simulations. The regression slopes *β* for the simulations were plotted against those for the observed data in Figure 4A, for 0-back condition effect, 2-back condition effect, and their differences respectively. Encouragingly, the simulated and actual *β* were nearly identical. Therefore, MINDy-X is able to generate realistic neural timeseries that recapitulate key statistics of the actual data (i.e., both the FC matrix and GLM response coefficients).

### 3.2 Topology of nontrivial brain dynamics across rest and task conditions

In previous sections, we verified that MINDy-X provides a plausible model for the generative process underlying observed brain activity across resting and task states. In the following sections, we further investigate this generative mechanism, by analyzing the attractor landscape of MINDy-X models. For each participant in the HCP dataset with sufficient low-motion neural data (N=537; see Methods), we ran simulations using their fitted MINDy-X model in the absence of noise, and with a constant input that was either *u* = (0, 0)^⊤^ (corresponding to resting state and task block interval), *u* = (1, 0)^⊤^ (0-back condition), or *u* = (0, 1)^⊤^ (2-back condition). We selected 120 random initial conditions and simulated fMRI data for at least 1600 time steps until the trajectories converged to the attractors. We found at least one stable equilibria or limit cycle in all models under all input conditions, and found no evidence of metastable or chaotic behavior. We visually examined the simulated trajectories of each model and excluded 14 (out of 537) models where the numerical attractor identification had potentially failed, and further excluded 12 participants with incomplete behavioral data or low performance (accuracy below 60% in 0-back or 2-back task), resulting in a sample size of 511 participants (models).

We analyzed the topology of the dynamical landscapes underlying the fitted models in terms of the attractors. The simulated trajectories from three models under resting state and 2-back task inputs are visualized in Fig 5A, along with their numerically identified attractors. The model displayed highly nontrivial attractor landscapes, including multistability, nonlinear oscillations and even multiple oscillatory orbits. To categorize the topology of dynamics, we counted the number of stable fixed points (FP) and limit cycles (LC) of each model under each input, as indicated by the title of each panel in Fig 5A. The distribution of the types of dynamics is displayed in Fig 5B, with color indicating the different rest/task states. Strikingly, during resting state, the models never showed monostable dynamics (1FP, 0LC). This finding was not due to a bias induced by fitting resting state and task fMRI data together, as the models that were fit on only resting state fMRI also never showed monostable dynamics (Supplementary Fig S1), indicating that methods that rely on the monostability assumption (e.g., linear system modeling) might miss important dynamical features that are present in rsfMRI data. In contrast, monostability dominated the dynamics during 0-back and 2-back conditions, suggesting a shift from a more flexible landscape during the resting state to a more rigid landscape under cognitive task demands. Despite such a global shift to monostability, the task state models still consisted of a sizable proportion of multistable (e.g., 2FP, 0LC) and oscillatory (e.g., 0FP, 1LC) dynamics, indicating that, just like resting state, the task state should also be considered as a non-linear dynamical system rather than monolithic state.

To better characterize the change in attractor landscape topology from resting to task state, we categorized the dynamics into three types: (1) oscillatory dynamics, with at least one stable limit cycle (LC *>* 1); (2) multistable dynamics, with no stable limit cycle but more than one stable equilibria (FP *>* 1, LC = 0); and (3) monostable dynamics, with no stable limit cycle and only one stable equilibria (1FP, 0LC). While such categorization is not based on strict topological equivalence between vector fields (which requires constructing a homeomorphism that is intractable for high-dimensional systems; but see our recent analytical developments in Chen et al., 2024), it summarizes the long-term behavior of the system: an oscillatory system can generate oscillations, while a monostable/multistable system will converge to one or several static states. We produced a contingency table for the type of dynamics each model displayed during resting state against those during 2-back task state (Fig 5C). A model was said to exhibit consistent (in contrast to bifurcated) dynamics between rest and 2-back states if the type of dynamics present was the same (in contrast to different) between the two states, as indicated by the two blocks with blue borders (in contrast to the four blocks with red borders) in Fig 5C. We found that in total, roughly 60% of models bifurcated, including roughly 53% that bifurcated to monostable dynamics, confirming a shift towards monostability under task drive.

Next, we examined the behavioral relevance of such topological change in neural dynamics. We compared the mean accuracy and log reaction time on 2-back task between participants whose model showed consistent (BIF-; blue blocks in Fig 5C) or bifurcated (BIF+; red blocks in Fig 5C) dynamics. Results were shown in Fig 5D. The BIF+ participants showed significantly higher accuracy, and, interestingly, also significantly slower reaction times, indicating a more cautious strategy (accuracy: Wilcoxon rank sum test *W* = 36433, *p* = 0.0021, rank biserial correlation *r* = 0.16, two-sample t-test *t*(509) = 2.9712, *p* = 0.0031, Cohen’s *d* = 0.27; log reaction time: Wilcoxon *W* = 34889, *p* = 0.0339, *r* = 0.11, t-test *t*(509) = 2.3499, *p* = 0.0192, *d* = 0.21). This effect might be driven by a trend to bifurcate towards monostability, as it was the participants who exhibited monostable dynamics in the 2-back condition that most clearly had higher accuracy and slower reaction times (Supplementary Figure S2).

### 3.3 Geometric proximity to task-related attractors predicts task performance

#### 3.3.1 Attractor geometry of fitted models

In the previous section, we showed that topological changes in dynamics were associated with higher task accuracy (but slower reaction time). Notably, in most cases, the dynamics bifurcated towards monostability during task state. We were next interested in examining the geometric properties of the attractor landscapes identified during task-states, and how these might be similar versus different from those present during resting states. Intuitively, the unique stable equilibrium preferentially observed in high-performing participants might be particularly relevant for task-related computations (Supplementary Figure S2). Therefore, we extracted the stable equilibria of all models during the resting state or with 2-back task state input. We obtained 750 equilibria from 333 models in resting state, and 689 equilibria from 465 models in the 2-back state (note that some “Oscillatory” models could also contain equilibria). Each equilibrium is a 200-dimensional vector in the state space (hidden space) of the model, representing a neural activation pattern over the 200 parcels (Schaefer et al., 2018). To understand their distribution over the population, we performed k-means clustering over the resting state equilibria and 2-back state equilibria separately. We used cosine distance for clustering to characterize the differences in activation patterns, but using Euclidean distance led to similar results. The optimal number of clusters *k* was selected between two to ten to obtain the most stable clustering solutions (Lange et al., 2004, see Supplementary Figure S3).

We visualized the attractors and cluster centroids in Fig 6A-D by linearly projecting them onto the first two “principal gradients” of whole-brain functional connectivity, defined in Margulies et al. (2016). The cortical parcel activation patterns corresponding to the cluster centroids were shown in the right panels. The resting state equilibria formed four well-defined clusters, characterized by the strong activation in default mode network (DMN), visual network (VIS), task-positive network (TPN), and salience network (SAL), respectively. The DMN (315 equilibria from 308 models) and TPN (319 equilibria from 311 models) clusters were much more popular than the VIS (56 equilibria from 51 models) and SAL (60 equilibria from 54 models) clusters. Importantly, not only the cluster centroids but also the individual equilibria showed modular activation patterns in line with functional brain networks (Yeo et al., 2011), suggesting that resting state functional networks might reflect non-trivial attractors (i.e., different from the mean activation pattern) present in nonlinear brain dynamics (Supplementary Figure S4).

In contrast to the resting state attractor dynamics (as well as the 0-back state, see Supplementary Figure S3), the 2-back state equilibria were found to be best described with a three-cluster solution. Interestingly, two of them resembled the DMN and TPN clusters in the resting state, while the other one showed a strong activation pattern located within the frontoparietal control network (FPN), similar to the learned MINDy-X input matrix and GLM regression slopes (Figure 3G). In fact, while less prominent than it was during the resting state, DMN was still the most popular cluster (299 equilibria from 280 models), followed by FPN (217 equilibria from 202 models) and TPN (173 equilibria from 165 models). Even within all monostable models, the DMN attractor (119 equilibria/models) was still slightly more popular than the FPN (113 equilibria/models) and TPN (37 equilibria/models) attractors. This finding suggests that apart from canonical trial-averaging neural patterns (i.e., the FPN “motif”), the presence of resting state attractor motifs are also present during cognitive task states and might have an important influence on task execution.

#### 3.3.2 Proximity to attractor motifs predict behavioral performance

Given the interesting coexistence of both resting-state-like and task-state-like attractors during the 2-back task blocks, it is natural to hypothesize that the geometry of the generative vector field might also predict individual differences in task performance. While there are many ways to characterize such geometry, here we explored two approaches that are conceptually similar to conventional methods, thus demonstrating the compatibility of the MINDy-X framework with existing methodologies. First, we conducted a state-based analysis similar to HMM approaches, which are often employed in time-varying functional connectivity (tvFC) studies (Lurie et al., 2020; Reinen et al., 2018). We extracted the parcel-level activation patterns (200-dimensional vectors) of each 2-back tfMRI volume, and labeled each volume by the closest (most similar) attractor cluster centroid (referred to as “motifs” from now on). For consistency with the methods used for clustering, here we utilized the cosine distance for labeling, although using Euclidean distance led to qualitatively similar results. We then calculated the proportion of volumes labeled by each motif for each participant, i.e., the proportion of time each participant spent in each motif state. An intuitive hypothesis is that spending more time in “task-relevant” motifs would be associated better task performance. Second, we computed the Euclidean distance between each volume and the motifs, again averaged across all 2-back volumes for each participant. Another intuitive hypothesis is that closer proximity to the task-relevant motifs would also be associated with better task performance.

We first present results that adopt an extreme group comparison approach, because of its greater analytic simplicity and transparency (Figure 6E-G). In the subsequent section, we follow-up this analysis with a formal GLM approach, to enable a more comprehensive characterization of individual differences across the full sample. We selected participants with either top 25% accuracy (“High” group, *n* = 128, Accuracy = 0.954 ± 0.0158, min = 0.938, max = 1) or bottom 25% accuracy (“Low” group, *n* = 138, Accuracy = 0.770 ± 0.0532, min = 0.625, max = 0.838). As expected, the high-performing group also had much faster log reaction time (Figure 6E).

We then analyzed whether the high and low-performing groups spent different amounts of time around each motif, akin to a HMM-type analysis of tvFC. We conducted a MANOVA predicting FPN and TPN state proportions (the DMN proportion was excluded, as it is defined to be equal to one minus the sum of the other two proportions) using participant group as the predictor variable. The MANOVA was significant: *F* (2, 263) = 4.7851, *p* = 0.009, Pillai’s Trace = 0.035. Post-hoc t-tests showed that DMN state proportion did not significantly differ across groups (High group: 31.5%; Low group: 32.3%), with *t*(264) = −1.0992, *p* = 0.273 (Holm-Bonferroni correction, same below), and Cohen’s *d* = −0.13 (CI: [−0.38, 0.11]). However, the FPN state proportion was significantly greater in high-performing participants (High group: 34.3%; Low group: 31.9%), with *t*(264) = 2.984, *p* = 0.009, and a small effect size of *d* = 0.37 (CI: [0.12, 0.61]). Correspondingly, the TPN state proportion dropped (High group: 34.2%; Low group: 35.7%), with *t*(264) = −2.295, *p* = 0.045, and a small effect size of *d* = −0.28 (CI: [−0.52, −0.04]). In summary, in line with the general scientific consensus regarding the role of FPN in cognitive control, we found that FPN state occupancy increased relative to the TPN state in the high-performing group, while the DMN state remained unchanged.

If state occupancy can predict differences in between high and low-performing participants, it also seems likely that the raw distances between neural activation patterns and attractor motifs may provide even richer information. Further, while the occupancy proportions in each state necessarily sums to one, the distances to the different motifs are relatively independent from each other. Consequently, we performed another MANOVA, this time predicting the mean distance to each motif, again using participant group as the predictor variable. This MANOVA was also significant: *F* (3, 262) = 7.837, *p <* 0.001, Pillai’s Trace = 0.082. Interestingly, post-hoc t-tests showed that the distances to the three motifs were all significantly reduced in the high-performing group, although the effect size was largest for the FPN motif. In particular, for the DMN motif, *t*(264) = −3.655, *p <* 0.001, a small effect size was observed *d* = −0.45 (CI: [−0.69, −0.20]). For FPN, *t*(264) = −4.658, *p <* 0.001, a medium effect size was observed *d* = −0.57 (CI: [−0.82, −0.33]). For TPN, *t*(264) = −3.360, *p <* 0.001, a small effect size was observed *d* = −0.41 (CI: [−0.65, −0.17]). This result appears to indicate that the DMN, FPN and TPN attractor motifs, treated as a set, may define a neural subspace that is suitable for 2-back task computation, with proximity to the FPN being potentially even more important than the other two.

### 3.4 Complementary role of neural geometry and topology in behavior prediction

Given that both the topology (e.g., rest-task bifurcation) and the geometry (e.g., mean distance to attractor motifs) of the generative vector fields seem to predict task performance, a natural follow-up question is to determine whether the two dimensions provide distinct or redundant information regarding individual differences in cognitive task performance. To probe this question, we fit GLMs to participant’s accuracy and log reaction time respectively, using combinations of the following predictors: (1) whether the model dynamics bifurcated from rest to task; (2) mean proportion of time in FPN and TPN states; and (3) mean distance to DMN/FPN/TPN motifs. We considered models with increasing complexity: (1) “Bifurcation”: only the bifurcation indicator; (2) “Occupancy”: only the state occupancy; (3) “Distance”: only the distances to motifs; (4) “Additive”: the bifurcation indicator as well as the distances to motifs; and (5) “Interaction”: the bifurcation indicator, motif distances, and the interaction between bifurcation and each distance predictor. While there are potentially many more predictors that can be extracted from MINDy-X, as well as many more reasonable combinations of predictors and their interactions, we selected the aforementioned models as typical examples for a MINDy-X-based exploratory analysis.

The model comparison results were summarized in Table 1, and the coefficients were presented in Table 2 and Table 3. As in Section 3.2, the bifurcation-only model significantly predicted both accuracy (*F* (1, 509) = 8.828, *p* = 0.003, adjusted *R*^2^ = 0.015) and reaction time (*F* (1, 509) = 5.522, *p* = 0.019, adjusted *R*^2^ = 0.008). On the contrary, the occupancy model only significantly predicted accuracy (*F* (2, 508) = 7.461, *p <* 0.001, adjusted *R*^2^ = 0.025), and not reaction time (*F* (2, 508) = 1.798, *p* = 0.167, adjusted *R*^2^ = 0.003). To predict accuracy, only the coefficient for FPN occupancy (0.194 ± 0.064, *t*(508) = 3.034, *p* = 0.0025) but not TPN occupancy (−0.018 ± 0.068, *t*(508) = −0.271, *p* = 0.787) was significant. As expected, in this model, a higher occupancy of the FPN state was associated with higher accuracy.

**Table 1:** Statistical models of behavior. Significant p-values (*<* 0.05) were shown in bold. “Adj. *R*^2^”: adjusted R squared. “vs. above”: F-test comparison between current model and the one above.

| Target | Model | $R^2$ | Adj. $R^2$ | $F$ | df | $p$ | AIC | BIC | vs. above |
| --- | --- | --- | --- | --- | --- | --- | --- | --- | --- |
| Accuracy | Bifurcation | 0.017 | 0.0151 | 8.83 | (1, 509) | **0.00311** | -1183. | -1170. |  |
|  | Occupancy | 0.0285 | 0.0247 | 7.46 | (2, 508) | **0.00064** | -1187. | -1170. |  |
|  | Distance | 0.0513 | 0.0456 | 9.13 | (3, 507) | **6.79e-06** | -1197. | -1176. |  |
| | Additive | 0.0582 | 0.0508 | 7.82 | (4, 506) | **4.02e-06** | -1199. | -1173. | $F = 3.76,$<br>$p = 0.053$ |
| | Interaction | 0.0681 | 0.0552 | 5.25 | (7, 503) | **8.35e-06** | -1198. | -1160. | $F = 1.78,$<br>$p = 0.149$ |
| log(RT) | Bifurcation | 0.0107 | 0.00879 | 5.52 | (1, 509) | **0.0192** | -552. | -539. |  |
|  | Occupancy | 0.00703 | 0.00312 | 1.8 | (2, 508) | 0.167 | -548. | -531. |  |
|  | Distance | 0.0224 | 0.0166 | 3.87 | (3, 507) | **0.00934** | -554. | -533. |  |
| | Additive | 0.0316 | 0.024 | 4.13 | (4, 506) | **0.00266** | -557. | -532. | $F = 4.86,$<br>$p = \mathbf{0.0279}$ |
| | Interaction | 0.0459 | 0.0326 | 3.45 | (7, 503) | **0.00127** | -559. | -520. | $F = 2.50,$<br>$p = 0.0585$ |

**Table 2:** Coefficients for models of accuracy. *B* and *β* indicate raw and standardized regression slopes, respectively. “s.e.”: standard error. “CI”: confidence interval.

| Model | Term | $B$ (s.e.) | $\beta$ | 95% CI of $\beta$ | $t$ | df | $p$ |
| --- | --- | --- | --- | --- | --- | --- | --- |
| Bifurcation | Bifurc | 0.0203 ($\pm 0.00683$) | 0.266 | [0.0901, 0.442] | 2.97 | 509 | **0.00311** |
| Occupancy | FPN time | 0.194 ($\pm 0.0641$) | 0.16 | [0.0566, 0.264] | 3.03 | 508 | **0.00254** |
| | TPN time | -0.0185 ($\pm 0.0684$) | -0.0143 | [-0.118, 0.0896] | -0.271 | | 0.787 |
| Distance | dDMN | 0.0224 ($\pm 0.0134$) | 0.154 | [-0.0267, 0.335] | 1.68 | 507 | 0.0945 |
| | dFPN | -0.0697 ($\pm 0.0202$) | -0.473 | [-0.742, -0.204] | -3.46 | | **0.00059** |
| | dTPN | 0.0253 ($\pm 0.0146$) | 0.155 | [-0.021, 0.332] | 1.73 | | 0.0841 |
| Additive | Bifurc | 0.0136 ($\pm 0.00703$) | 0.178 | [-0.00272, 0.359] | 1.94 | 506 | 0.0535 |
| | dDMN | 0.0156 ($\pm 0.0138$) | 0.108 | [-0.079, 0.294] | 1.13 | | 0.258 |
| | dFPN | -0.0579 ($\pm 0.021$) | -0.393 | [-0.674, -0.113] | -2.75 | | **0.00608** |
| | dTPN | 0.0199 ($\pm 0.0148$) | 0.122 | [-0.057, 0.301] | 1.34 | | 0.181 |
| Interaction | Bifurc | 0.587 ($\pm 0.272$) | 0.17 | [-0.0129, 0.352] | 2.16 | 503 | **0.0312** |
| | dDMN | 0.0449 ($\pm 0.0236$) | 0.309 | [-0.0107, 0.629] | 1.9 | | 0.0581 |
| | dFPN | -0.0777 ($\pm 0.0359$) | -0.527 | [-1.01, -0.0484] | -2.16 | | **0.031** |
| | dTPN | 0.0322 ($\pm 0.0256$) | 0.198 | [-0.112, 0.507] | 1.26 | | 0.21 |
| | Bifurc:dDMN | -0.0429 ($\pm 0.0291$) | -0.295 | [-0.689, 0.0985] | -1.47 | | 0.141 |
| | Bifurc:dFPN | 0.0295 ($\pm 0.0443$) | 0.2 | [-0.39, 0.791] | 0.667 | | 0.505 |
| | Bifurc:dTPN | -0.0203 ($\pm 0.0314$) | -0.125 | [-0.504, 0.254] | -0.647 | | 0.518 |

**Table 3:** Coefficients for models of log reaction time.

| Model | Term | $B$ (s.e.) | $\beta$ | 95% CI of $\beta$ | $t$ | df | $p$ |
| --- | --- | --- | --- | --- | --- | --- | --- |
| Bifurcation | Bifurc | 0.0298 ($\pm 0.0127$) | 0.211 | [0.0346, 0.387] | 2.35 | 509 | **0.0192** |
| Occupancy | FPN time | 0.202 ($\pm 0.12$) | 0.0904 | [-0.0147, 0.195] | 1.69 | 508 | 0.0916 |
| | TPN time | 0.212 ($\pm 0.128$) | 0.0888 | [-0.0162, 0.194] | 1.66 | | 0.0972 |
| Distance | dDMN | 0.0634 ($\pm 0.0251$) | 0.236 | [0.0526, 0.42] | 2.53 | 507 | **0.0118** |
| | dFPN | -0.0356 ($\pm 0.0378$) | -0.131 | [-0.404, 0.142] | -0.94 | | 0.348 |
| | dTPN | 0.00774 ($\pm 0.0274$) | 0.0257 | [-0.153, 0.205] | 0.283 | | 0.778 |
| Additive | Bifurc | 0.0289 ($\pm 0.0132$) | 0.205 | [0.0216, 0.388] | 2.2 | 506 | **0.0286** |
| | dDMN | 0.049 ($\pm 0.0259$) | 0.183 | [-0.00671, 0.372] | 1.9 | | 0.0587 |
| | dFPN | -0.0104 ($\pm 0.0394$) | -0.0383 | [-0.323, 0.246] | -0.265 | | 0.791 |
| | dTPN | -0.00377 ($\pm 0.0278$) | -0.0125 | [-0.194, 0.169] | -0.136 | | 0.892 |
| Interaction | Bifurc | 0.282 ($\pm 0.508$) | 0.239 | [0.0543, 0.424] | 0.556 | 503 | 0.579 |
| | dDMN | 0.0012 ($\pm 0.0442$) | 0.00446 | [-0.319, 0.328] | 0.0271 | | 0.978 |
| | dFPN | 0.113 ($\pm 0.0671$) | 0.415 | [-0.0698, 0.9] | 1.68 | | 0.0932 |
| | dTPN | -0.0645 ($\pm 0.0479$) | -0.214 | [-0.527, 0.0988] | -1.34 | | 0.179 |
| | Bifurc:dDMN | 0.0758 ($\pm 0.0544$) | 0.283 | [-0.116, 0.681] | 1.39 | | 0.164 |
| | Bifurc:dFPN | -0.189 ($\pm 0.0827$) | -0.693 | [-1.29, -0.0961] | -2.28 | | **0.023** |
| | Bifurc:dTPN | 0.088 ($\pm 0.0587$) | 0.292 | [-0.091, 0.676] | 1.5 | | 0.135 |

**Table 4:**
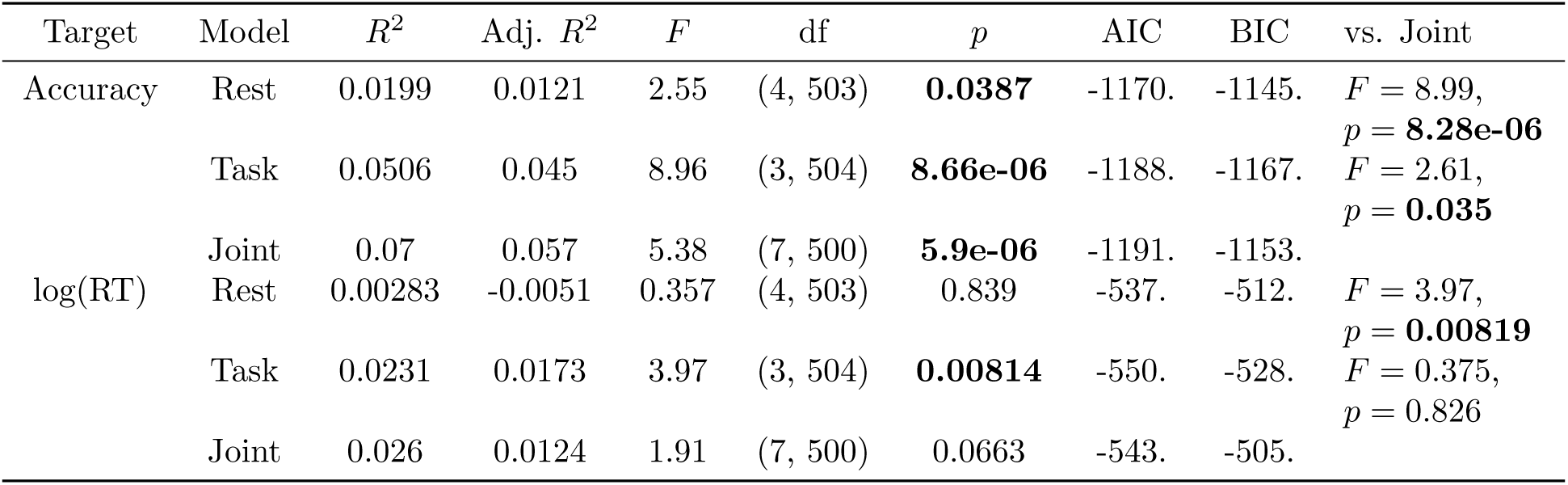
Comparison of resting state and task-based models.

The distance model was significant for both accuracy (*F* (3, 507) = 9.131, *p <* 0.001, adjusted *R*^2^ = 0.046) and reaction time (*F* (3, 507) = 3.871, *p* = 0.009, adjusted *R*^2^ = 0.017). For accuracy, only the coefficient for distance to the FPN motif was negative and significant (−0.0697 ± 0.020, *t*(507) = −3.458, *p <* 0.001), while the other two were positive and marginally significant (*p <* 0.1). For reaction time, only the distance coefficient for DMN was positive and significant (0.063 ± 0.025, *t*(507) = 2.527, *p* = 0.01), while the other two were not significant. In short, in this model, closer proximity to the FPN motif predicted higher accuracy, while closer proximity to the DMN motif predicted faster reaction time.

The additive model showed a similar picture. The model explained significantly more variance than the bifurcation-only model for both accuracy (adjusted *R*^2^ = 0.051, F-test *F* (3, 507) = 7.376, *p <* 0.001) and reaction time (adjusted *R*^2^ = 0.024, *F* (3, 506) = 3.638, *p* = 0.013). The model also explained marginally significantly more variance than the distance model for accuracy (*F* (1, 506) = 3.762, *p* = 0.053), and significantly more for reaction time (*F* (1, 506) = 4.863, *p* = 0.028). For accuracy, the bifurcation effect remained positive and marginally significant (0.0136 ± 0.007, *t*(506) = 1.935, *p* = 0.053), and the FPN distance effect remained negative and significant (−0.058 ± 0.021, *t*(506) = −2.755, *p* = 0.006). For reaction time, the bifurcation effect remained positive and significant (0.0289 ± 0.013, *t*(506) = 2.196, *p* = 0.029), and the DMN distance effect remained positive and marginally significant (0.049 ± 0.026, *t*(506) = 1.895, *p* = 0.059).

Based on the additive model, both the topological and geometric features provided unique contributions to behavioral prediction. However, the geometric features themselves also differed significantly across bifurcated and consistent models (MANOVA for all three distance measures against bifurcation group, *F* (3, 507) = 16.613, *p <* 0.001, Pillai’s Trace = 0.0895), leading to an interesting question about the contribution of their interaction. Therefore, we proceeded to the interaction model. The interaction model explained marginally significantly more variance than the additive model only for reaction time (adjusted *R*^2^ = 0.0326, F-test *F* (3, 503) = 2.503, *p* = 0.059), but not for accuracy (adjusted *R*^2^ = 0.055, *F* (3, 503) = 1.783, *p* = 0.149). Interestingly, in the interaction model for reaction time, the main effect of bifurcation and distance to the DMN motif were no longer significant. Instead, the main effect of distance to the FPN motif was positive and marginally significant (0.113 ± 0.067, *t*(503) = 1.682, *p* = 0.0932), while the interaction with bifurcation was negative and significant (−0.189 ± 0.083, *t*(506) = −2.281, *p* = 0.023). Therefore, according to this model, for participants without a rest-task bifurcation, closer proximity to the FPN motif predicted faster responses (0.113 ± 0.067), while for participants who did exhibit bifurcation, FPN proximity predicted slower responses instead (−0.076 ± 0.048). Critically, both the topological and geometric features contributed to the prediction of individual differences in behavior, and they provided both synergistic and unique information about the accuracy and speed of task computation.

### 3.5 Unique predictive power of resting state and task state neural geometry

In the previous sections, we demonstrated that the neural geometry present during 2-back task state can predict key aspects of individual differences in task performance. Given that a key assumption of the MINDy-X framework is that the resting state and task state neural dynamics are generated by a same unifying mechanism, it is intriguing to further investigate whether the dynamical features of attractor geometry during the resting state by itself can also predict task performance; and further, if the joint resting and task state dynamics predict performance in a unique or redundant manner, relative to when one or the other is examined in isolation.

To this end, we calculated the mean distances between observed neural states and the four resting state attractor motifs (DMN, VIS, TPN, and SAL) based on every participant’s rsfMRI data. To rule out the leakage of information between sessions during model training, we re-defined the resting state motifs based on MINDy models trained only on rsfMRI, although the redefined motifs were nearly identical to the ones shown in Figure 6. We retained a subset of 508 participants (models) after removing re-trained rsfMRI models with numerical issues. We then predicted the 2-back task accuracy and log reaction time of participants, using the four distance measures defined on resting state fMRI, similar to the “Distance” model in the previous section, but now only including resting-state data and attractor motifs. The rsfMRI measures failed to predict log reaction time (*F* (4, 503) = 0.357, *p* = 0.8392, adjusted *R*^2^ *<* 0). However, they significantly predicted accuracy (*F* (4, 503) = 2.547, *p* = 0.039, adjusted *R*^2^ = 0.012). The estimates were negative and significant for distance to DMN (−0.277 ± 0.116, *t*(503) = −2.388, *p* = 0.017) and SAL (−0.649 ± 0.261, *t*(503) = −2.484, *p* = 0.013). The estimate were positive and significant for distance to VIS (0.609 ± 0.303, *t*(503) = 2.010, *p* = 0.045), and not significant for TPN (0.301 ± 0.188, *t*(503) = 1.596, *p* = 0.111). Namely, fluctuation around DMN and SAL motifs (and away from the VIS motif) during the resting state was associated with better performance during N-back cognitive task states.

We then analyzed whether the resting state distance measures explained unique or overlapping variance in accuracy with the task state distance measures. Interestingly, the correlations between rsfMRI and tfMRI distance measures across individuals were quite weak (|*ρ*| *<* 0.1), suggesting that they might hold unique predictive power. On the new subset of participants, we re-fit the task state “Distance” model, which explained around 4.5% of variance (*F* (3, 504) = 8.955, *p <* 0.001, adjusted *R*^2^ = 0.045). We then fit another model to predict accuracy using the four distance measures from rsfMRI and the three distance measures from tfMRI, which explained around 5.7% of variance (*F* (7, 500) = 5.378, *p <* 0.001, adjusted *R*^2^ = 0.057), roughly the same as the sum of explained variance from the resting-state-only (1.2%) and task-state-only (4.5%) models. As expected, the combined model explained significantly more variance than using tfMRI alone (*F* (4, 500) = 2.609, *p* = 0.035). Therefore, resting state and task-state geometrical features were not simply linearly related; instead, they had unique predictive power in accounting for cognitive individual differences.

## 4 Discussion

In this article, we tested the hypothesis that fMRI-measured whole-brain dynamics during resting states and controlled task states are generated by a unifying dynamical mechanism. Towards this aim, we introduced the MINDy-X framework for joint modeling and analysis of rsfMRI and tfMRI timeseries. We validated that in the HCP dataset, MINDy-X provided a better fit to both rsfMRI and N-back tfMRI data, relative to linear models. Further, we demonstrated that MINDy-X can generate realistic simulations that recreate the correlational structure (FC) and steady state response (GLM slopes) of the original data. Meanwhile, MINDy-X model parameters were also interpretable and test-retest reliable within individuals. We then applied MINDy-X to analyze the changes in attractor landscapes across rest and N-back working memory task states. Topologically, the models possessed a variety of nontrivial dynamics ranging from monostability and multistability, to oscillations. The task effect manifested as a predominant trend, observed in the majority of participants, to bifurcate towards monostable dynamics under task conditions. Further, this trend was associated with higher accuracy and more cautious responding. Geometrically, the models showed nontrivial attractors that clustered well according to functional brain network activations. The proximity between individual brain activation patterns and model-defined attractor motifs, uniquely present in task and resting states, jointly predicted individual differences in task performance. More generally, we provided novel and rigorous evidence for a unifying neurocomputational mechanism underlying whole-brain dynamics across multiple cognitive states. We also developed a new modeling and analysis framework that can be used to reveal the interesting complementary roles of neural topology and geometry in cognitive processing, opening a wide range of new possibilities for studying the neural mechanisms of cognition in individuals and across different population groups.

### 4.1 Unifying mechanisms for whole-brain dynamics across rest and task states

Until recent years, resting-state and task-based functional neuroimaging research have been developing quite independently from each other. Early efforts for joint analysis of rsfMRI and tfMRI showed that the statistics of brain dynamics such as FC are highly consistent between resting and task states (Cole et al., 2014; Krienen et al., 2014), but also that the distinction between the two may uniquely encode cognitive individual differences (Schultz & Cole, 2016). To further understand the role of resting state dynamics in cognition, it is necessary to advance from statistical associations to mechanistic explanations. Apart from summarizing the data, mechanistic models embed hypotheses regarding the generative process that gives rise to the observed data (Friston et al., 2019), offering explanatory value. Further, mechanistic models generate verifiable predictions regarding perturbations of the system, offering predictive value (Bassett et al., 2018). Such predictions provide theoretical guidance to experimental interventions, such as exogenous neurostimulation (Singh, Cole, et al., 2022), and consequently, could form a rigorous computational foundation for translational research.

Prior to our study, there has been significant progress in the mechanistic modeling of fMRI, and, to a certain degree, joint modeling of rsfMRI and tfMRI. Here we briefly review three major types of such models: activity flow models, Hidden Markov Models (HMMs), and (continuous-state) dynamical system models. Note that there also exist black-box machine learning models (Yang et al., 2025) and single-neuron biophysical models for fMRI (Lu et al., 2024), but we restrict our focus on gray-box models which strike a balance between descriptive validity and mechanistic interpretability.

Activity flow models are feedforward models where FC guides the transformation between regional neural representations in order to implement cognitive computation (Ito et al., 2020). Activity flow models aim to capture the spatial dimension of neural computation, but they place less emphasis on the temporal dimension. On the contrary, HMMs capture the temporal dynamics of whole-brain activity in terms of probabilistic switching between discrete states (Vidaurre et al., 2018). These states are usually identified by whole-brain activation patterns or time-varying FC patterns (Vidaurre et al., 2025), but in principle, they could also be the parameters of a lower-level generative model (Schwamb et al., 2026). The HMM neural states can be associated with different mental states such as spontaneous thoughts (Kucyi, 2018), attention or arousal fluctuations (Nakuci et al., 2025; Song et al., 2023), and emotions (Kragel et al., 2022). Dynamical-system models like MINDy-X extend the discrete states implemented in HMMs to a continuous state space characterization (Friston et al., 2019; Kashyap et al., 2025; Mitjans et al., 2023; Sanz Leon et al., 2013; Sip et al., 2023; Tanner et al., 2024; Vahidi et al., 2024). In dynamical system models, the temporal structure is embedded in the topology and geometry of the generative vector field, while the spatial structure is given by the mapping between the anatomical space and the model’s state space. Although dynamical system models can arguably provide the most comprehensive mechanistic description of brain activation patterns, the computational relevance of such a description has been almost completely unexplored prior to this study.

In this paper, we developed a new modeling framework MINDy-X, which addresses several methodological challenges associated with the joint modeling of rsfMRI-tfMRI data: (1) the ability to capture fundamentally nonlinear phenomenon, such as multistability and limit cycles; (2) sensitivity to individual differences; (3) scalability to whole-brain characterization; (4) flexibility to include various block-level and event-related task effects; and (5) mechanistic interpretability. While there has been significant progress in related literatures (Englert et al., 2025; Kashyap et al., 2025; Vahidi et al., 2024), according to our knowledge, MINDy-X is the first modeling framework to achieve all these goals simultaneously. By implementing such a framework, the use of MINDy-X allowed us to demonstrate for the first time that the topology and geometry of a unified whole-brain attractor landscape is associated with individual differences in cognitive task performance, thus providing a new perspective on the cognitive brain.

### 4.2 Analyzing the unified dynamics

In this study, we demonstrated that whole-brain dynamics can be modeled as a unified dynamical system, consisting of interconnected neural masses, and modulated by cognitive state and individual variation. We next discuss how this mechanism can be studied at two different levels: the level of underlying components, and the level of emergent dynamics.

First, this unified whole-brain neural mechanism can be studied in terms of its lower-level components. When adopting the MINDy-X framework, such components are the parameters of the model, which can be analyzed akin to conventional FC and GLM techniques. MINDy-X parameters can be interpreted in simple terms: effective connectivity weights *W* indicate causal influence from one region to another; the reciprocal of slope *α* encodes the variability of neural activity within each region; decay rates *D* reflect regional excitation-inhibition balance; and input weights *B* encode the effect of experimental contrasts on each region’s neural activity. Taking advantage of both resting state and task fMRI to obtain parameter estimates, MINDy-X can potentially provide more reliable metrics of these parameters at the individual level. In fact, the connectivity measures were quite consistent across resting state and “task residual” (subtracting GLM-estimated task effects) fMRI data (Cole et al., 2014). When combining both modalities, the estimated connectivity measure also predict individual cognitive traits better (Easley et al., 2023). Similarly, task effect estimates might also benefit from “subtracting” resting state dynamics from the task activation data (Kashyap et al., 2025; Singh, Wang, et al., 2022). Apart from providing more reliable estimates, MINDy-X extends the estimate of effective connectivity (EC) to that occurring between several hundred interacting neural populations, which is a significant expansion from that possible using alternative approaches, such as dynamic causal modeling, which due to computational tractability is limited to estimating connectivity from much fewer regions (Friston et al., 2019). This expansion has the potential to enable whole-brain graph-theory based EC analyses of the type that could previously only be conducted on structural connectivity (SC) or functional connectivity (FC) estimates (Bullmore & Sporns, 2009). Such analyses enable investigation of the cause-effect structure and information-flow patterns present across the whole brain, potentially bridging the gap between SC and FC approaches (Suárez et al., 2020).

Second, and of potentially greater interest, with MINDy-X it becomes possible to analyze unifying mechanisms at the emergent dynamics level; a level which is not accessible through FC- or GLM-type approaches. When adopting a continuous-state dynamical system framework such as MINDy-X, the whole-brain activity can be viewed as reflecting noise-driven trajectories that traverses an attractor landscape (Deco, Jirsa, & McIntosh, 2013). Importantly, at this level, the attractor landscape becomes the primary mechanistic substrate for cognitive phenomena, while the underlying components that produce such landscapes serve as secondary mechanisms (Barack & Krakauer, 2021). As exemplified in our current analyses of the HCP dataset and N-back task, the effects of cognitive states and individual differences can be characterized in terms of modulations to the topology and geometry of attractor landscapes, and not just in terms of model parameters. MINDy-X is particularly suitable for such study as here it revealed a richer set of nonlinear landscapes compared to previous methods. Most critically, MINDy-X explicitly models the effect of experimental contrasts on the generative dynamics, which can provide a new vocabulary for understanding and characterizing such experimental manipulations at a mechanistic level. As a concrete illustration of this point, within the current MINDy-X framework, cognitive effects can be viewed in two ways. A sustained effect, such as a block or condition effect, can be considered as one that primarily modulates the dynamical landscape, which potentially involves changing its topology. Conversely, a transient or event-related effect can be considered as a driving force of activation that that may help the brain navigate the geometry of this landscape.

Of course, it is crucial to demonstrate how dynamics-level explanations provide new scientific insights into the brain. Whole-brain level dynamical systems analysis, such as those done with MINDy-X, are less likely to be helpful for questions involving localized neural activity, e.g., representation of visual stimulus within primary visual cortex. However, many interesting cognitive phenomena have been associated with network-level or whole-brain-level neural dynamics. For example, mind-wandering and spontaneous thoughts are believed to involve the dynamical interactions between DMN, FPN and dorsal/ventral attentional networks (Christoff et al., 2016; Kucyi, 2018). By combining MINDy-X modeling with thought sampling procedures (Gonzalez-Castillo et al., 2021), one can probe the link between neural attractors and dimensions of thought contents (Christoff et al., 2016). Similarly, practices and training interventions that may impact mind-wandering and spontaneous thought patterns, such as mindfulness meditation, have also been characterized in terms of network-wide interactions and dynamical processes (Czajko et al., 2024; Dagnino et al., 2024; Lin et al., 2026; Vohryzek et al., 2025). Thus, MINDy-X models may be a critical tool for understanding mindfulness-related brain states and how these modulate attractor dynamics present during both spontaneous thought and under conditions that may otherwise promote mind-wandering. Along another line of research, (Kragel et al., 2022) suggests that the spontaneous dynamics of emotions might be best characterized in terms an HMM switching between several emotional brain states, leading to an interesting question about the emotional relevance of MINDy-X attractor motifs.

The MINDy-X architecture is also particularly suitable for studying cognitive control. While one should not equate the “control input” in MINDy-X with the “control signal” in the brain (Medaglia, 2019), MINDy-X does provide an efficient way to model both the sustained and transient effect of control demands on brain dynamical landscape. Therefore, MINDy-X could be particularly instrumental for characterizing distinct control processes or modes, such as those postulated in the Dual Mechanisms of Control framework (DMCC; Braver, 2012). In particular, a tantalizing possibility is that proactive and reactive control might differentially reflect modulations in the topology and geometry of the whole-brain dynamics, respectively. Last but not least, attractor-based analysis has already shown value in research on sustained attention. As a recent example, Song et al. (2025) showed that the relationship between ongoing neural states and MINDy attractors were associated with moment-to-moment fluctuations in attention, suggesting that attention was encoded in the attractor landscapes of whole-brain dynamics.

### 4.3 Future directions

By adopting the new perspective of a unifying dynamical mechanism underlying both resting state and task-related activation patterns, profound shifts could potentially occur in the way that cognitive and clinical neuroscience research is conducted, including advances in both the analytic methods and experimental designs that are commonly employed in research investigation.

In terms of mechanistic modeling, there are many directions in which the MINDy-X framework could be extended and utilized. Currently, the influence of experimental contrasts is modeled by an intercept term in the dynamics, which additively modulates the activity of each region. While this formulation is a good starting point, it is also somewhat limited in terms of modeling task-state modulations. In fact, it seems quite plausible to assume that cognitive state variations will not only modulate regional activity, but also change the effective connectivity between regions (Friston et al., 2003). Indeed, prior studies have shown that connectivity can change systematically across cognitive tasks (Cole et al., 2013). Extending MINDy-X to include task-based modulation of connectivity weights would be a valuable direction to explore. Another possible improvement is a better measurement model that includes regional and inter-subject variability in the HRF kernel (Aguirre et al., 1998). While it is quite common to assume a fixed measurement model in whole-brain analysis of fMRI data (Sip et al., 2023), regional variability in HRF might lead to subtle mis-estimates that could complicate interpretation of connectivity weights (Seth et al., 2015). Methods already exist to incorporate HRF variation within the MINDy resting-state framework (Singh, Wang, et al., 2020), so although adapting these methods to MINDy-X will increase the complexity of parameter estimation, there is a clear-cut path for how to overcome this limitation of current modeling.

While modeling frameworks like MINDy-X capture the nonlinear dynamics of whole-brain activity, new analytic methods are now needed to more effectively interpret the modeled dynamics. One of the most interesting directions, as shown in this paper, is to compare the neural topology and geometry observed across different cognitive states. However, currently, such analyses are limited to be relatively coarse-grained in nature. For example, we categorized the topology of dynamics into merely three types (monostability, multistability, oscillation). This is because rigorous evaluation of topological congruency is extremely difficult within high-dimensional nonlinear models like MINDy-X. Advanced machine-learning-based techniques could be valuable for comparing and reasoning about these models (Chen et al., 2024). Another interesting direction is to analyze non-linear dynamical systems models, such as MINDy-X, using principles from well-established frameworks such as control theory. Within such frameworks, the cognitive effects can be viewed as “control inputs” to the unifying dynamics. Since such dynamics exhibit fundamentally nonlinear phenomena like multistability and limit cycles, traditional linearization-based analysis would likely fall short (Gu et al., 2015). Reassuringly, new methods have been recently developed to analyze and control such dynamics, which do not rely on linearization approaches (Tamekue & Ching, 2025; Tamekue et al., 2025, 2026).

Further, the unification of resting state and task-performing brain dynamics also suggests new experimental designs. The theory of unifying brain dynamics is particularly synergistic with the paradigm of naturalistic cognitive tasks involving continuously changing inputs (such as movie-watching) and behavioral measures (Huk et al., 2018). Such paradigms allows researchers to directly link the generative dynamics of the brain to the generative dynamics that give rise the behavioral states (Raut et al., 2025), thus establishing a new route to understand their causal relationship. More radically, if we treat cognitive state variations as modulations to the unifying dynamics, with a proper model of such dynamics (e.g., a MINDy-X model), we can design tasks and stimuli that might be optimized to induce certain brain dynamics. For example, we might be able to design a task for a given individual to consistently engage their frontoparietal control network (FPN) as an attractor for cognitive training purposes, instead of forcing such activations, which is the current state-of-the-art in neurostimulation research.

Last but not least, while we only demonstrate the value of MINDy-X modeling and analysis of unifying dynamics in healthy young adults, it is clearly valuable to extend this approach to development, aging, and patient populations. Many studies have tried to understand and predict aging or neuropsychiatric disorders using dynamical models of rsfMRI (e.g., Garćes de Marcilla Lappin et al., 2026; Nakagawa et al., 2013), and, very recently, both rsfMRI and tfMRI (Englert et al., 2025; Pine et al., 2025). However, the task-induced reconfiguration of unifying brain dynamics and how this might differ across distinct populations is a question that has remained largely unexplored. Compared to the rsfMRI-only approach, the joint modeling method might have substantially increased power and sensitivity to more directly link changes in brain dynamics with variations in cognitive task performance.

Overall, we believe that the language of dynamical systems theory can not only help clarify the neural mechanisms of cognition, but it also may potentially pave the way towards a more naturalistic and comprehensive cognitive neuroscience research paradigm. One that can fully enable rigorous and ecologically valid scientific study of the rich, complex repertoire of behaviors engaged in by the human species.

## Acknowledgments

We thank Geoffrey Goodhill and Janine Bijsterbosch for their helpful comments. Part of this work was funded by R21MH132240 from National Institutes of Health and N00014-22-S-F0 from Multidisciplinary University Research Initiatives (MURI) Program.

## Supplementary Figures

**Figure S1:**
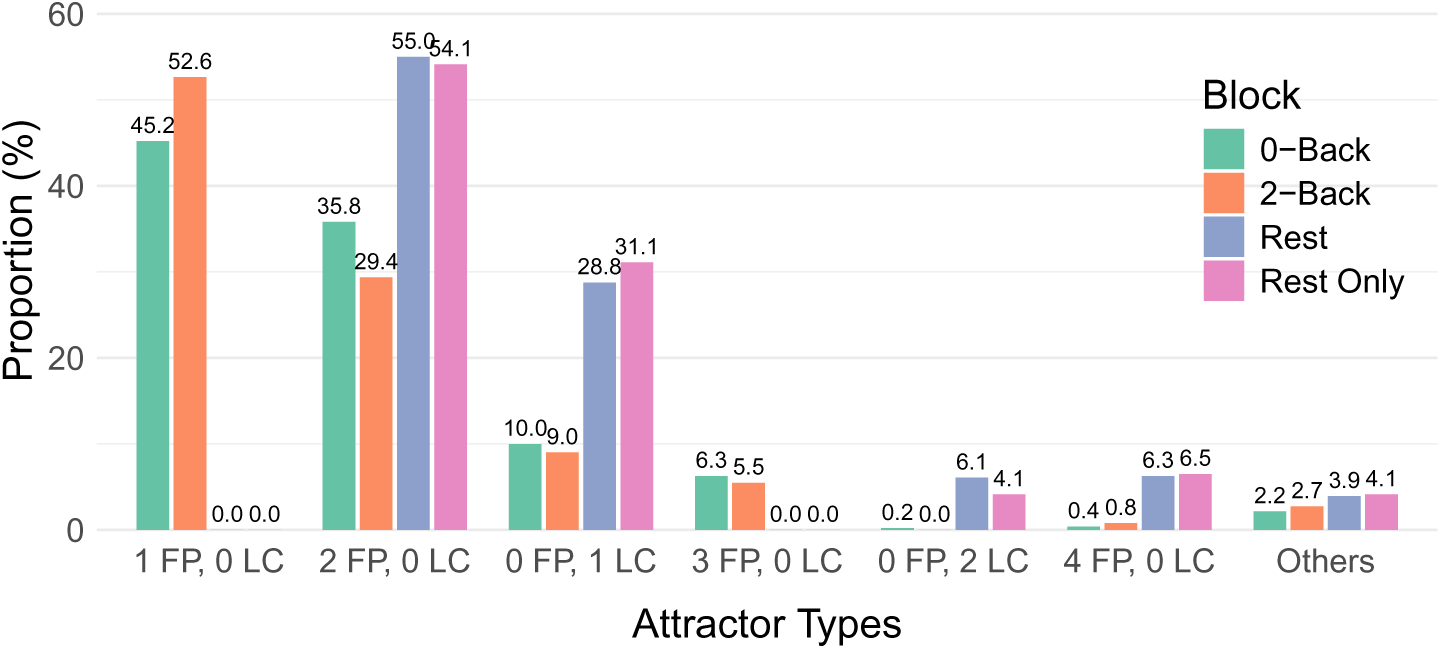
Taxonomy of dynamics across resting-state only models and joint models in each condition. The resting-state only models were used in the analysis in Section 3.5.

**Figure S2:**
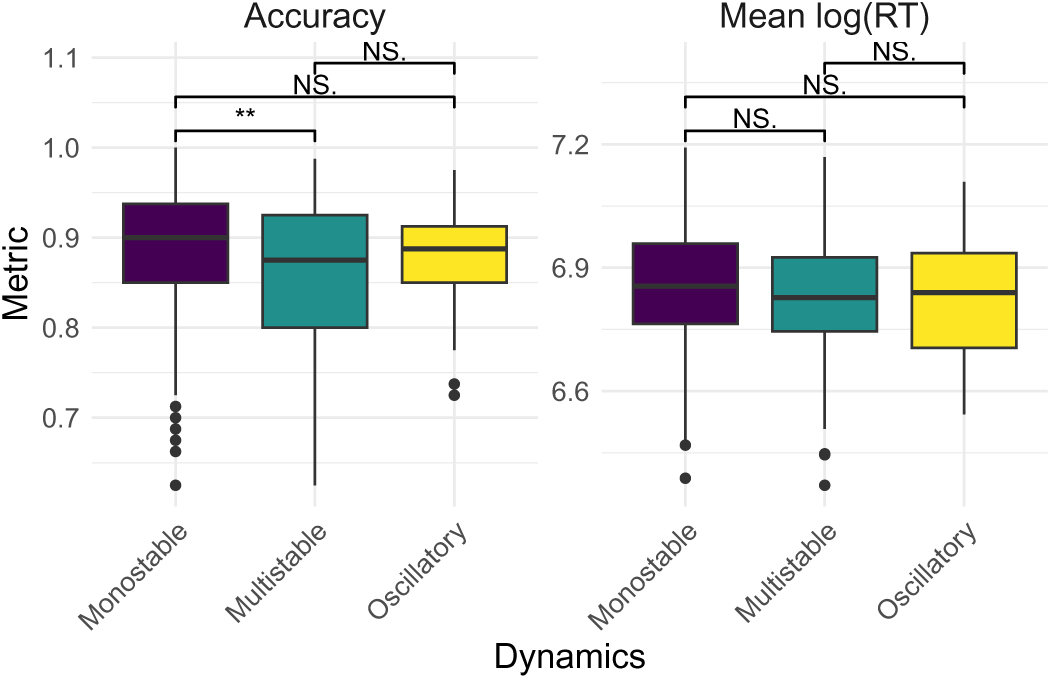
Behavioral differences across participants with different dynamics in 2-back condition. ANOVA was significant for accuracy (*F* (2, 508) = 5.507, *p* = 0.00431) and marginally significant for log reaction time (*F* (2, 508) = 2.601, *p* = 0.0752). Only the differences between monostable and multi-stable groups for accuracy was significant (Tukey’s HSD, estimate = 0.0237, CI: [0.0067, 0.0407], adjusted *p* = 0.0032). “N.S.”: not significant.

**Figure S3:**
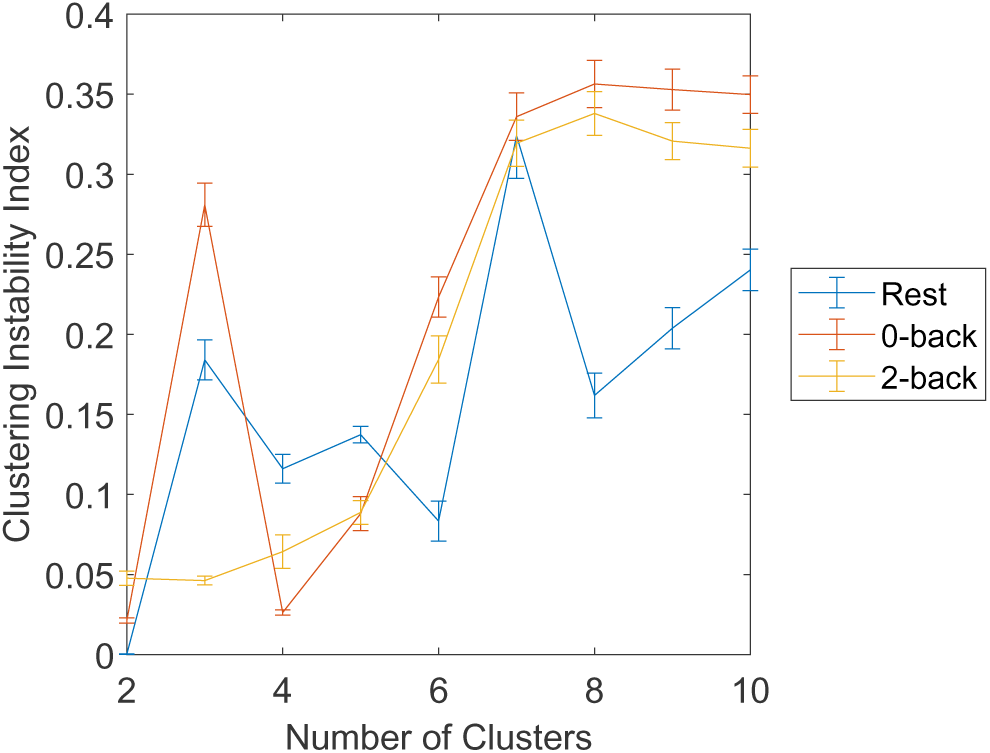
Clustering instability for attractors in each condition.

**Figure S4:**
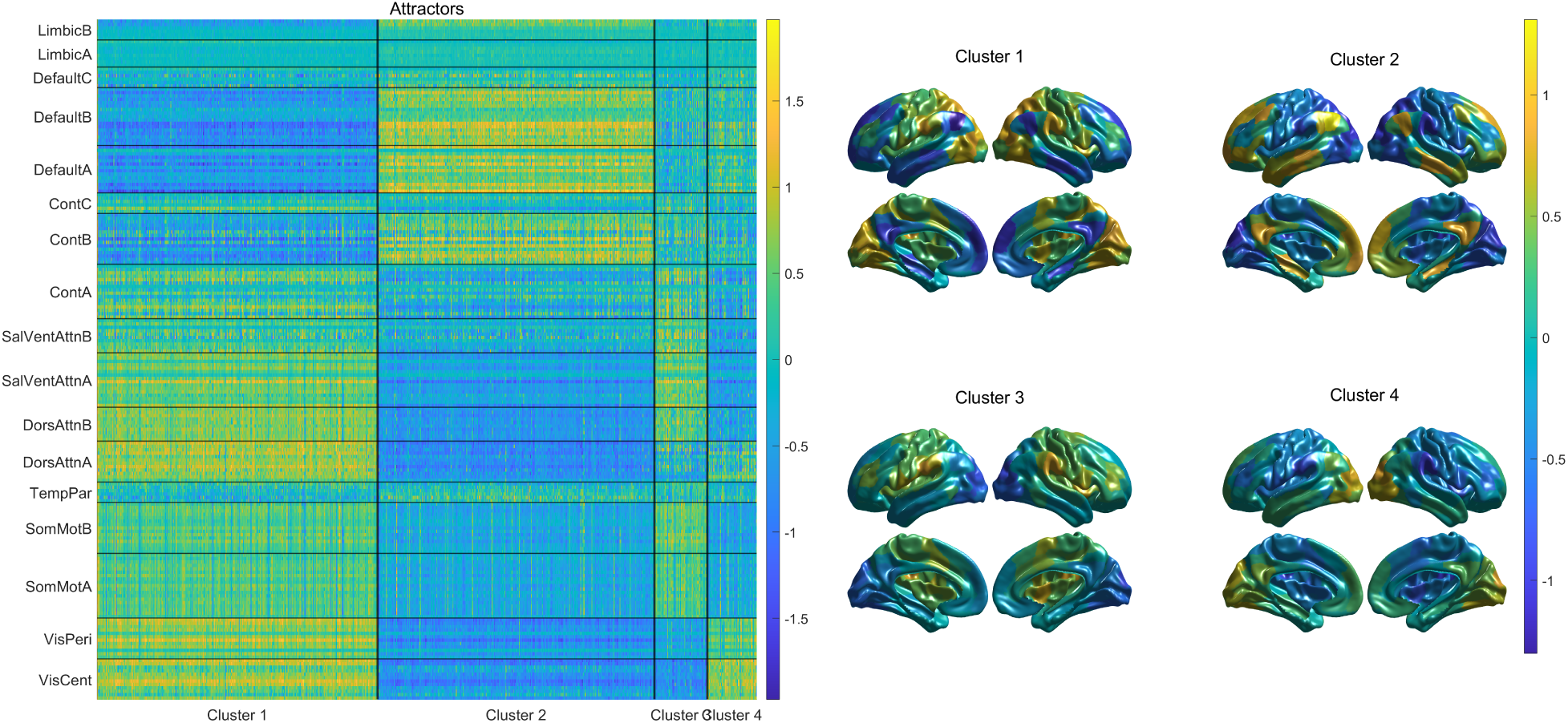
Clustering resting state attractors. Left: all resting state attractors. Each row represents one region and each column represents one attractor. Color indicates the coefficient (activation). Right: cluster centroids visualized as activation pattern over the cortex.

**Figure S5:**
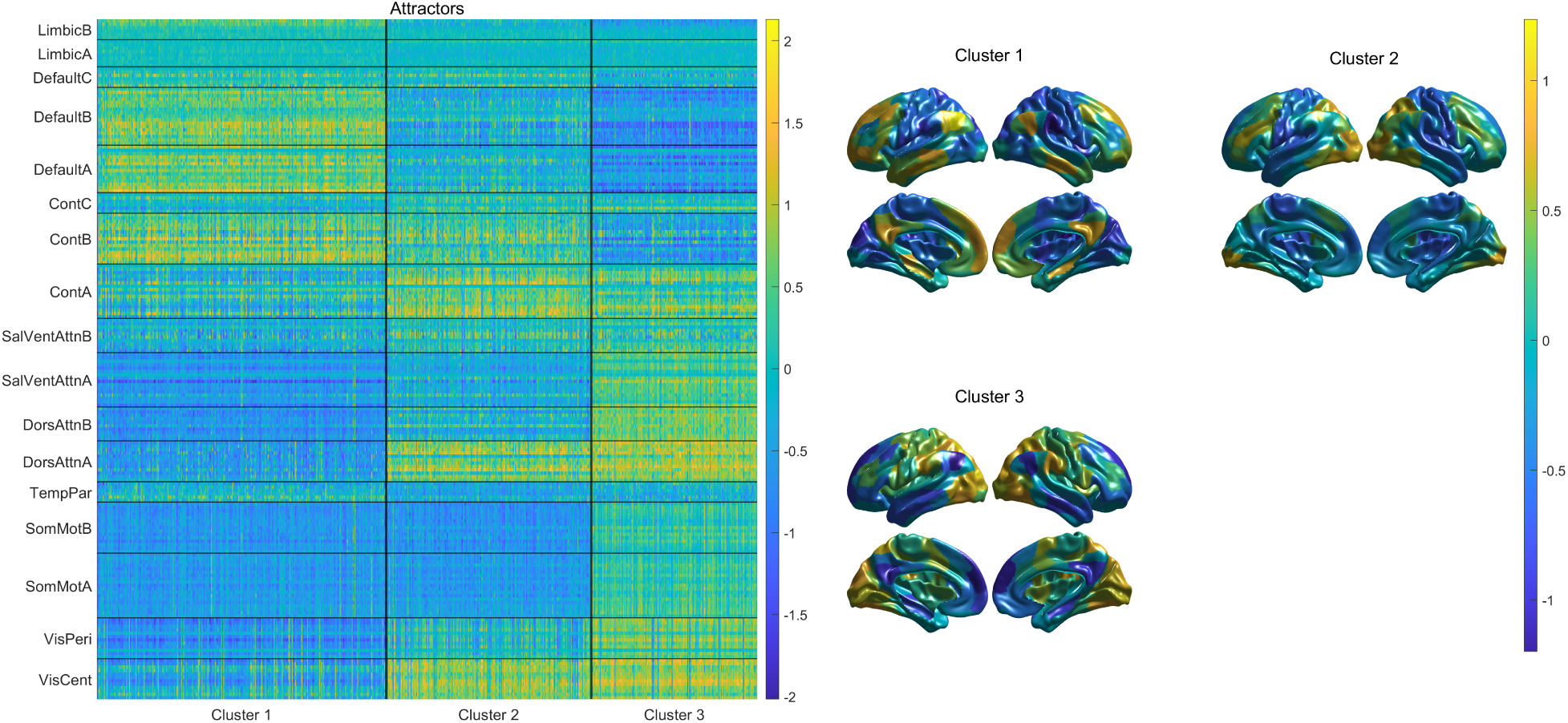
Clustering 2-back condition attractors.

